# Linking cross-species hemodynamics with neuronal activity in prefrontal cortex during abstract sequence viewing

**DOI:** 10.64898/2026.10.23.761183

**Authors:** Katherine E Conen, Brent W Stewart, M Vanessa Rivera Núñez, Hannah Hyde, Nadira Yusif Rodriguez, Aarit Ahuja, Debaleena Basu, Theresa H McKim, Theresa M Desrochers

## Abstract

Humans and other primates have a remarkable ability to identify and predict abstract patterns that evolve through time, but the neural mechanisms that support this capability remain poorly understood. Lateral prefrontal cortex (LPFC) has been implicated in abstract sequence tracking in both humans^1–5^ and macaques^5–9^, but it is unclear whether underlying processes are shared across species. Further, most existing evidence relies on hemodynamic responses, which can have variable relationships with neuronal activity^10–14^. We leveraged a combination of human fMRI, monkey fMRI, and electrophysiology to bridge these gaps. We found that humans and macaques show functionally homologous hemodynamic activation in LPFC during abstract sequence viewing, and neuronal responses within fMRI-targeted regions in macaques contain information about abstract and sensory variables in parallel. However, neuronal activity did not directly mirror hemodynamic responses, and abstract sequence responses were found both near and far from fMRI peaks. In contrast, distinct dynamics of spiking activity were selectively present near fMRI peaks, suggesting that temporal coordination across neurons may influence hemodynamic activity in prefrontal cortex. Together, these experiments establish a link between human brain activity, macaque fMRI, and single neuron responses, providing a window into the neural processes that mediate abstract sequence detection.

## Main

Abstract pattern recognition is a fundamental building block of complex cognition. An extraordinary range of human behavior, from language and music to cooking, weaving, and computer programming, depends on our capacity to extract higher-order rules from sequences of events and apply them flexibly across new contexts. One tractable exemplar of this capacity is the detection of abstract (or algebraic) sequences: structured patterns defined not by specific stimuli or motor actions, but by the relationships between elements (e.g., AAAB, ^^^&, TTT9), which unfold over time in a series of discrete steps^15,16^. Behavioral studies have shown that both humans and monkeys learn abstract sequence rules and generalize them to new stimuli^17–21^. Even in the absence of task demands, abstract sequence deviants evoke changes in hemodynamic activity, gaze behavior, and pupillary response^5,6,9,21^, suggesting that abstract sequence tracking may be an innate component of cognition in these species. However, few studies have examined the correlates of abstract sequence detection at the neuronal level^8,9,18^. The relationship between hemodynamic signals and neuronal spiking remains unclear, particularly in regions of prefrontal cortex that have been linked to abstract cognition^10,11,14,22–25^.

We combined human fMRI, macaque fMRI, and macaque electrophysiology to identify the neural correlates of abstract sequence tracking across species and methodologies (Fig.1A). We focused primarily on lateral prefrontal cortex (LPFC), which has been linked to abstract sequence processing across a range of converging methodologies, including neuroimaging^1,4–6,18^, electrophysiology^8,9,26^, TMS^1^, and lesion studies^27–29^. Our approach consisted of four components: (1) identifying deviant-responsive subregions of LPFC in both species using fMRI; (2) directly correlating hemodynamic responses across species to test for functional homology; (3) recording extracellular spiking activity from fMRI-defined regions to characterize abstract sequence tracking at the neuronal level; and (4) identifying how responses at the neuronal level vary with distance from fMRI-defined peaks.

**Figure 1.**
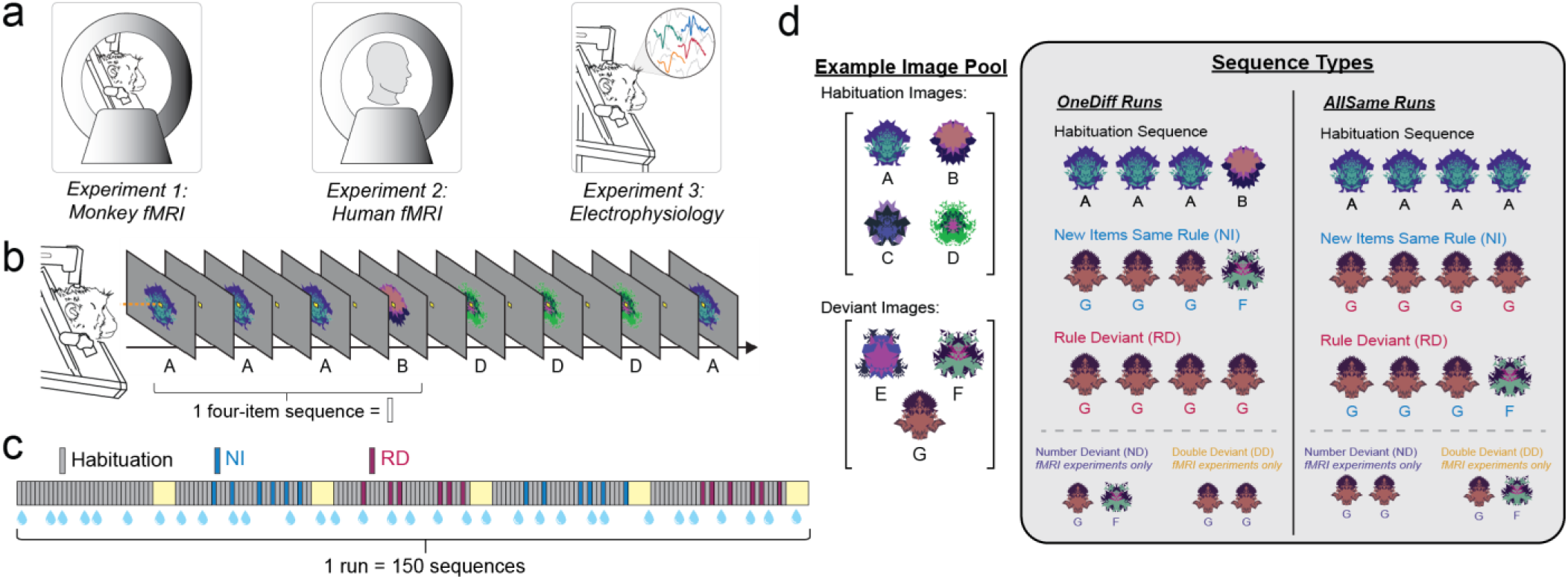
Cross-species analysis of brain activity using a no-report abstract sequence viewing task. **(A)** Responses to abstract sequence deviants were studied using a no-report sequence task in three experiments: monkey fMRI, human fMRI, and monkey electrophysiology. **(B)** Example of two habituation trials from a run with the OneDiff sequence pattern. **(C)** Schematic of the run structure used in monkey electrophysiology experiments. Each bar represents one four-item sequence. Gray bars represent habituation sequences, while blue and purple bars represent New Image (NI) and Rule Deviant (RD) sequences respectively. Yellow squares indicate 14s inter-block fixation periods. Droplets represent juice reward, which was delivered on a graduated schedule contingent on good fixation, independent of stimulus and sequence timing. **(D) Left:** Examples of the habituation and deviant image pool from one run. **Right:** Illustration of the habituation and deviant sequence types for the two run patterns.

### Humans and macaques show similar hemodynamic responses during no-report sequence viewing

If humans and macaques rely on similar mechanisms for abstract sequence tracking, related brain regions should be activated across species. In a previous study, we identified a specific subregion of macaque LPFC that showed increased hemodynamic responses when sequences violated an expected abstract rule^6,7^. This area, the posterior fundus of area 46 (p46f), is in an anatomically similar position to the human dorsolateral prefrontal cortex (DLPFC)^30,31^ and shows similar functional connectivity patterns as the human rostrolateral prefrontal cortex (RLPFC)^32^, which has been causally linked to abstract sequence tracking in other tasks^1,3,4^. These similarities suggest that human LPFC might play a comparable role to monkey p46f during abstract sequence viewing. To test this hypothesis, we collected fMRI data from human participants during the same no-report abstract sequence task run in monkeys (see Methods; Fig.1B-D, Supplementary Fig.1B). Briefly, participants maintained fixation on a central fixation spot while a series of fractal images was displayed during fMRI scanning. Images were displayed in groups of four images that either followed the habituated standard pattern (e.g., three same, one different, AAAB), or, rarely, deviated by using new images (new images with same rule, NI) or by using a different rule (rule deviant, RD). To control for the frequency of images and isolate changes in the abstract sequence rule, we compared hemodynamic responses on RD and NI sequences. These responses served as an index for areas of the brain that were tracking abstract sequence information.

Abstract sequence responses in human LPFC paralleled the monkey results. When modeling each sequence at the onset (see Methods), we found bilateral clusters in the lateral prefrontal cortex (LPFC) showing significantly greater activation for RD>NI (Fig. 2A; Supplementary Table 1). The left LPFC cluster included a peak in the inferior frontal gyrus (MNI *xyz* = −36, 12, 22), while the right LPFC cluster included peaks in the inferior and middle frontal gyrus (MNI *xyz* = 40, 18, 20; 54, 30, 18; 32, 24, 34) and in the precentral gyrus (MNI *xyz* = 46, 4, 28). Both clusters extended beyond the inferior frontal gyrus into more dorsal frontal regions. The left cluster also extended into the medial frontal cortex, including the supplementary motor area (SMA), pre-SMA, and middle cingulate gyrus. Notably, the right LPFC cluster overlaps with a cluster observed during a previous active sequence task^2^ and adjoins a more anterior RLPFC cluster from that same study. Further, the general distribution of areas that respond to RD>NI were similar in humans and monkeys across the frontal cortex beyond the LPFC, with both species also showing activation in medial prefrontal cortex and dorsal premotor cortex (Fig.2A, Supplementary Tables 1-2). In sum, abstract sequence deviants drive a similar constellation of activation in both humans and macaques, even when these deviants are not relevant to task performance or reward.

**Figure 2.**
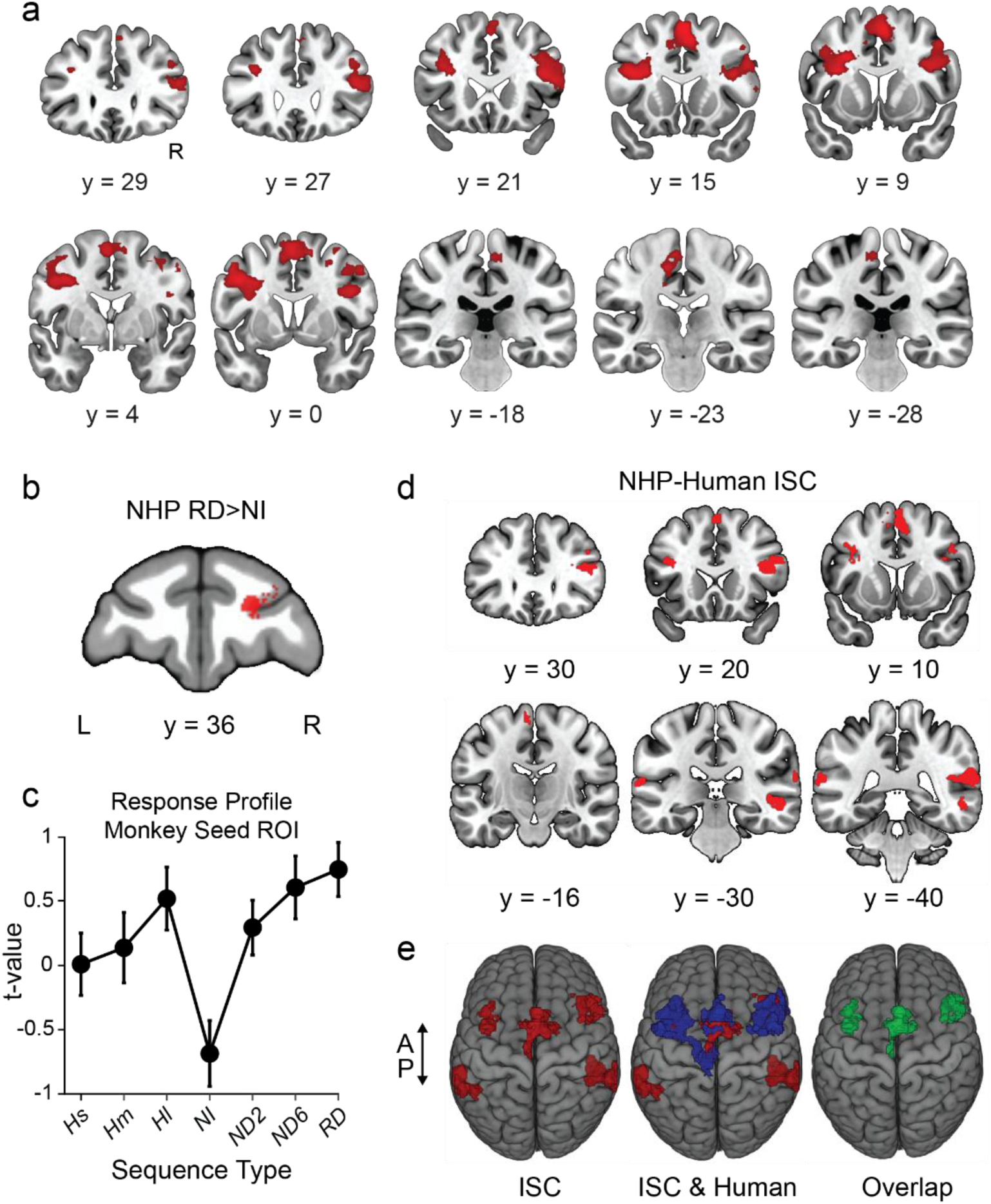
Similar hemodynamic responses to abstract sequence deviants in humans and macaques. **(A)** Coronal slices show the group-level RD>NI contrast from the onsets model (N = 28), overlaid on the MNI152 template. Clusters were defined using a height threshold of p < .005 (uncorrected) and an extent threshold of k = 1,327 voxels, and survived cluster-level FWE correction (p < .05). The left hemisphere is displayed on the left. **(B)** Subregion in right p46f (Rp46f) of monkeys that responds more strongly to RD than NI sequences used to define the seed ROI for ISC analysis. **(C)** Activation profile in the Rp46f seed region from monkeys across three habituation and four deviant sequence types (Hs, Hm, Hl = short, medium, and long-duration habituation sequences; ND2 = 2-image number deviants, ND6 = 6-image number deviants). Error bars show ±SEM. **(D)** Human brain regions exhibiting responses across conditions significantly correlated with monkey Rp46f (FWEc < 0.05, height p < 0.005 unc., extent = 239). **(E)** Rendered brains displaying monkey Rp46f-to-human ISC (red), monkey Rp46f-to-human ISC with the human RD>NI contrast from (a) (blue), and their overlap (green).

### Hemodynamic responses to abstract sequences are directly correlated across species

We next asked whether responses in these regions were functionally homologous across all task conditions. To test this question, we performed interspecies correlation (ISC)^33,34^ between macaque and human fMRI data, directly comparing activation patterns. For ISC, we applied a vector of average hemodynamic responses across seven conditions (NI, RD, 2- and 6-image number deviants, and three different habituation timings) in a region of interest (ROI) to correlate the profile of t-values, voxel by voxel, across the brain of the same or different species, producing a “map” of areas that show the same pattern of responses across conditions (see Methods). Within-species ISC in monkeys and humans was used to validate the method, with both species showing a high degree of overlap between ISC maps and univariate activations (Supplementary Fig.2, Supplementary Tables 3-4). We then performed monkey-to-human ISC using the region identified by the monkey RD>NI contrast in right p46f as a seed ROI (Fig.2BC). This analysis revealed significant ISC clusters bilaterally in the human LPFC, superior temporal cortex, and supplementary motor area (Fig.2D, Supplementary Table 5). Clusters of significant ISC overlapped with the previously observed human RD>NI contrast in bilateral LPFC and supplementary motor area (Fig.2E; 44% of ISC voxels overlapped with human RD>NI contrast). Human-to-monkey ISC using a right LPFC seed region from the human RD>NI contrast similarly revealed significant ISC in monkey right DLPFC, adjacent to the univariate cluster but not directly overlapping it (Supplementary Fig.2, Supplementary Table 6). Together, these results indicate that hemodynamic responses show distinct cross-species homology in LPFC areas that track sequential information.

### Neuronal responses differentiate sequence deviants but do not mirror hemodynamics

Our fMRI results showed that abstract sequence processing recruits functionally homologous brain regions in humans and macaques, but the precise nature of neuronal responses within these regions remained an open question. Bulk hemodynamic signals may obscure important features of activity in a network. Moreover, while studies from sensory regions have found correlations between hemodynamic signals and neuronal spiking^13,35–37^, the relationship is relatively weak and may not be consistent across different brain regions and task contexts^10,14,22–25^. To address this gap, we performed subject-specific, fMRI-targeted recording in macaque LPFC, analyzing the activity of 887 units from two monkeys (Monkey B: n = 469 units, Monkey W: n = 418 units; Fig.3A, Supplementary Fig.3).

**Figure 3.**
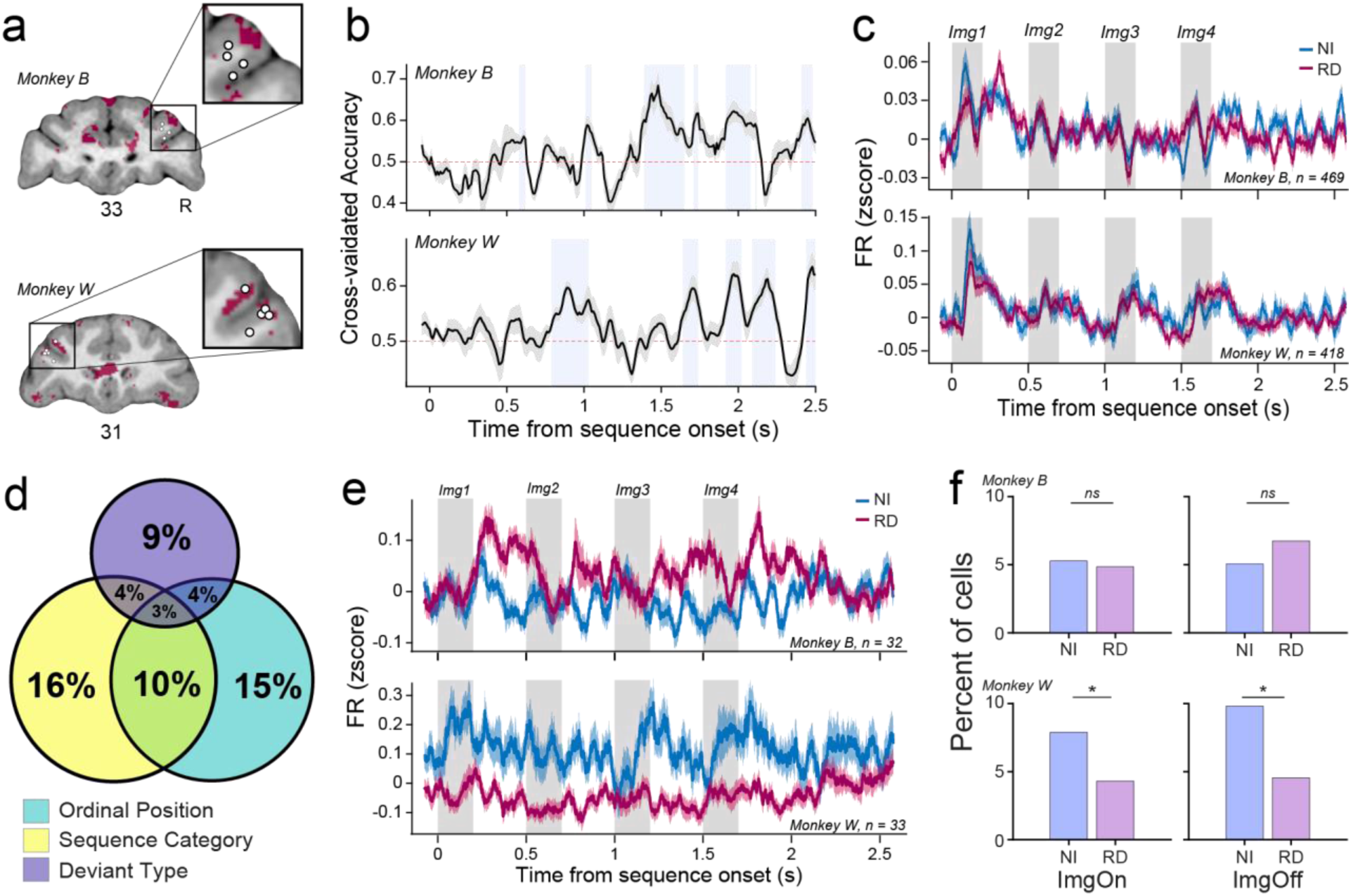
Neuronal responses to abstract sequence variables in functionally targeted subregions of LPFC. **(A)** Example fMRI-guided recording sites from monkey B (top) and W (bottom), displayed on a T1-weighted anatomical scan collected from the same monkey. White circles show the location of the deepest electrode site on a given recording day. Overlays show fMRI contrasts for RD>NI onsets (maroon) (p < 0.05, uncorrected). Insets show a magnified view of target recording sites on the example slice. The inset for monkey W is mirrored for visual comparison to monkey B. **(B)** Accuracy of deviant type decoding on held-out data across the course of the sequence for monkey B (top) and monkey W (bottom). Traces show the mean ± SEM across ten train/test splits. Blue shaded rectangles indicate intervals with significant above-chance decoding (p < 0.05 in at least two contiguous bins, Benjamini-Hochberg FDR-corrected one-sided t-test). Red dotted line shows 50% decoding accuracy (chance). Decoding was conducted using LDA on sliding firing rates sampled with 200ms bin size and 10ms step size from 50ms before the sequence start to 2.5s after the sequence end. X-axis values are aligned to the middle of the time bin. Peak decoding accuracy was 0.68 for monkey B and 0.64 for monkey W. **(C)** Mean responses to RD and NI sequences in monkey B (top) and W (bottom) across all neurons. Responses to the two deviant types were not significantly different in either the ImgOn or ImgOff time windows (one-way ANOVA; monkey B: ImgOn, p = 0.35, F = 0.88; ImgOff, p = 0.68, F = 0.17; monkey W: ImgOn, p = 0.23, F = 1.47; ImgOff, p = 0.17, F = 1.89; degrees of freedom = 1 for all). Shaded rectangles correspond to the fractal presentation time. Traces show mean ± SEM. **(D)** Proportion of units with a significant response to the three main task variables, pooled across monkeys and time windows. **(E)** Deviant type responses show differences across multiple sequence positions. Top: Mean response across units from Monkey B with significantly higher response to RD>NI sequences in the ImgOff time window. Bottom: Mean response across units from monkey W with a significantly higher response to NI>RD in the ImgOn time window. Firing rates were z-scored by run and by cell before averaging and were sampled with 50ms sliding time bins (1ms step size). Traces show mean ± SEM. **(F)** Proportion of significant RD>NI and NI>RD responses for each monkey and time window. Asterisks indicate a significant difference between RD>NI and NI>RD proportions (two-sided exact binomial test, monkey B: ImgOn, p = 0.89, ImgOff, p = 0.50; monkey W: ImgOn, p = 0.049, ImgOff, p = 0.013).

First, we asked whether neuronal responses in LPFC directly mirrored the RD>NI hemodynamic signals observed in this task. A decoding analysis found that deviant type could be categorized with above-chance accuracy on held-out data in both monkeys, indicating that there were systematic differences in population activity on NI and RD trials (Fig.3B). Significant decoding was primarily observed during and after the fourth image in the sequence, when NI and RD sequences diverged. Briefer windows of significant decoding also occurred around position 2 of the sequence, potentially reflecting contextual expectations about deviant type in the current block. Notably, decoding performance did not arise from overall differences in firing rate on RD and NI trials, and neither monkey showed a significant difference in their mean response to NI and RD sequences (Fig.3C). These results suggest that hemodynamic activity does not directly parallel firing rates in LPFC, but may reflect distinct patterns of activity across neurons, potentially revealing differences in the network state.

The abstract sequence viewing task involves a combination of perceptual variables (stimulus identity, sequence timing) and more abstract features (habituation vs. deviant category, deviant type), which may both evoke responses in LPFC neurons. To examine this possibility in more detail, we analyzed responses to three primary task variables: ordinal position (1-4), sequence category (habituation vs. deviant), and deviant type (NI vs. RD). We also performed secondary analyses of image selectivity separately for deviant and habituation trials. Firing rates were measured 50-350ms after image onset and offset (“ImgOn” and “ImgOff” time windows). Responses in 60% of cells showed a significant relationship with at least one task variable, and approximately half of task-sensitive neurons responded to multiple variables (Fig.3D, Supplementary Fig.4, Supplementary Table 7). Effects of ordinal position and sequence category were prevalent in both monkeys, and were frequently found in the same neurons. These responses often appeared to track salient visual features of the task: most position responses were highest at the sequence start (Pos1), and sequence category responses often showed selectivity for image-identity in addition to sequence category (Supplementary Fig.5AB). Responses to deviant type were also present in both monkeys, though they were slightly less common (Fig.3DE). One monkey had a greater fraction of RD>NI responses, while the other showed the opposite pattern (Fig.3F). Interestingly, while neurons rarely showed significant interactions between deviant type and other task variables (Supplementary Fig.5EF), the magnitude of deviant type responses was greater for the neuron’s preferred stimulus (Supplementary Fig.5GH). These results demonstrate that stimulus and timing-related variables are represented in parallel with abstract, higher-order features such as deviant type in LPFC, even when they are not required for task performance.

**Figure 4.**
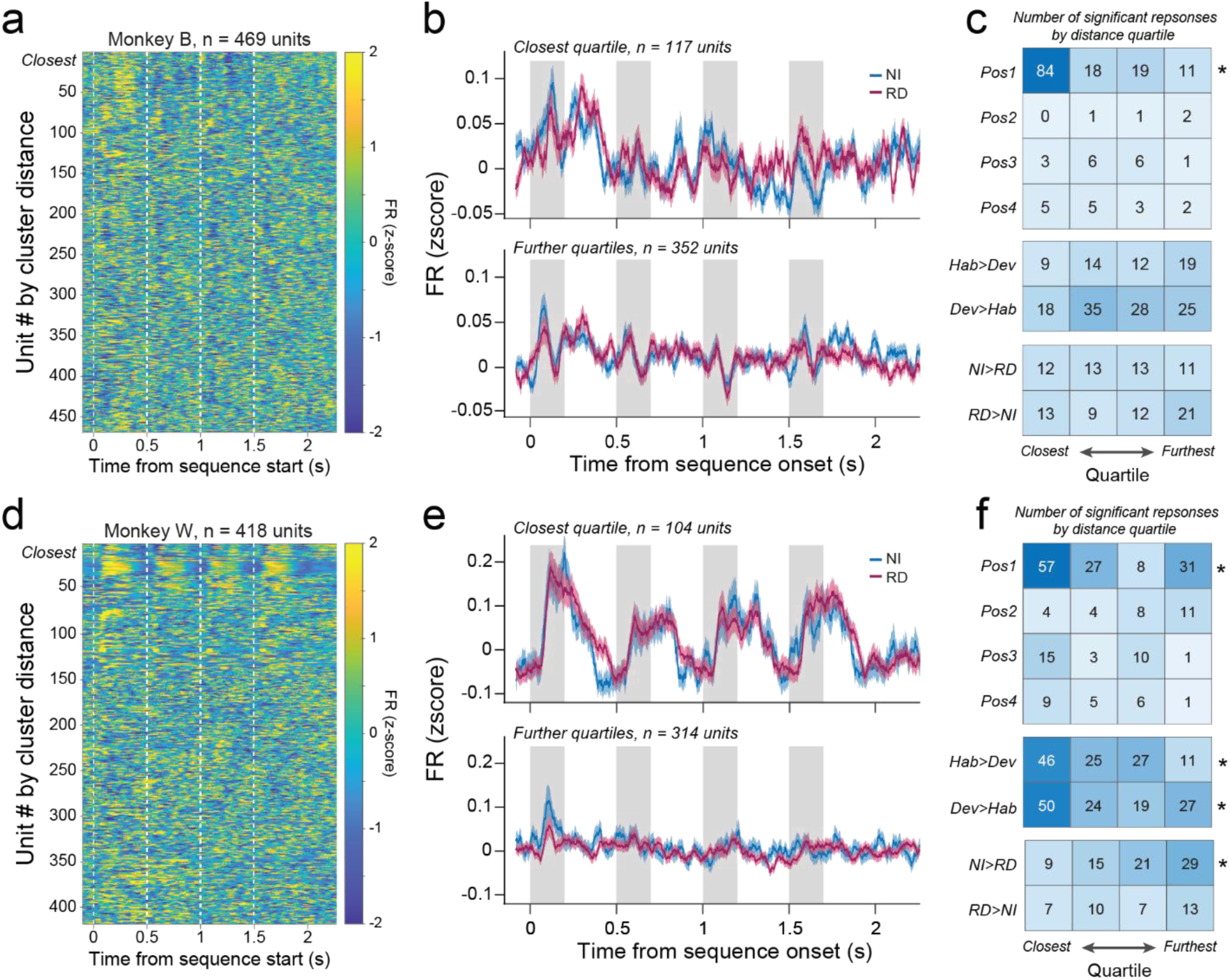
Neuronal response profiles vary with distance from hemodynamic peak. **(A,D)** Firing rates of each unit during deviant sequences, sorted by distance from the RD>NI hemodynamic peak. Firing rates were calculated using 50ms sliding time bins with 1ms step size. Vertical dotted lines indicate the onset of each image in the sequence. **(B,E)** Mean response to NI and RD sequences in the quartile of units closest to the cluster peak and the other three quartiles from Monkey B (B) and Monkey W (E). Shaded regions correspond to fractal display time. Traces show mean ± SEM. **(C,F)** Number of units with a significant effect of each task variable for Monkey B (C) and Monkey W (F) in each distance quartile, pooled across time windows. Asterisks indicate a significant effect of cluster distance (logistic regression, p<0.05 for both coefficient and model). Panels (A-C): Monkey B; panels (D-F): Monkey W.

Neuronal responses related to hemodynamic activity may be more common closer to fMRI-defined cluster peaks. To investigate this question, we identified the largest RD>NI cluster in the recorded hemisphere for each monkey and found the coordinates for the highest peak in that cluster (Supplementary Table 8). Units at different distances from the cluster peak showed qualitatively distinct response dynamics (Fig.4ABDE, Supplementary Fig.6-7). Using logistic regression, we tested whether distance between the recording site and the cluster peak affected the probability of finding task-related variables. Contrary to our expectations, RD>NI responses were distributed broadly throughout LPFC and were not preferentially found near the cluster peak, although monkey W had a higher proportion of NI>RD responses further from the cluster peak (Fig.4CF, Supplementary Fig.8AB, Supplementary Tables 9-10). Decoding contribution indices also did not correlate with distance from the cluster center (Monkey B: *r* = 0.039, *p* = 0.41; Monkey W: r = 0.045, p = 0.48; Spearman correlation on estimated Shapley values^41,42^). In contrast, we observed strong effects of distance for position-related responses: neurons that responded most strongly to position 1 were heavily concentrated near the RD>NI onsets cluster in both monkeys, and the mean firing rate near this cluster showed a clear phasic peak at the sequence start (Fig.4BE, Supplementary Fig.8). Thus, while hemodynamic peak locations do reflect spatial heterogeneity in neuronal responses, they are primarily correlated with phasic sequence start responses rather than responses to deviant type. In conjunction with our decoding results, these findings suggest that hemodynamic responses in LPFC have a complex relationship with neuronal activity, reflecting response timing, network coordination, and potentially subthreshold patterns of activity across neurons rather than mean spiking activity in a specific cortical region.

## Conclusion

These experiments establish the first direct throughline between human brain activity, monkey fMRI, and single cell neuronal responses in the same monkeys during an abstract sequence task. Interspecies correlation analyses identified functionally homologous regions in LPFC of monkeys and humans, suggesting that components of abstract sequence tracking may be evolutionarily conserved across primates. In humans, these areas adjoin and overlap with regions of dorsal and rostral LPFC that have been linked to abstract sequence tracking in active tasks^1,4^, and may be part of a broader prefrontal network involved in abstract sequence processing.

While both hemodynamic and neuronal responses differentiated between NI and RD sequences, neuronal responses did not mirror the RD>NI signal observed in fMRI. These results contrast with previous work in sensory regions, where positive correlations between neuronal activity and hemodynamic responses have been observed^13,35–37^. Several factors could account for these differences, including differences in cortical structure across regions^38^, the degree of feedforward processing^12,24,39^, the distribution of neurotransmitters^25,40–42^, and the no-report structure of the task^42,43^. Notably, several studies have found small or negative correlations between hemodynamics and spiking in certain contexts, suggesting that these relationships vary substantially depending on brain region and task structure^12,22,24,44^. Considered together, these results suggest that fMRI responses and neuronal spiking may represent complementary methods for understanding network activity, rather than similar signals on a different spatiotemporal scale. In particular, while spiking can provide more direct information about information processing within a region, fMRI may reflect a combination of network-level activity patterns, changes in inter-regional communication, specific cell types, and non-neuronal contributions^14,39,45–48^. Understanding the distinct physiological processes that contribute to hemodynamic signals will expand our ability to understand brain function in health and disease.

Across both species and methodologies, our results show that LPFC distinguishes between new sequences that match a familiar pattern and those that deviate from it, even when this distinction is not necessary for task performance. These results add to a growing body of evidence that LPFC tracks features of abstract sequences^1–3,5–9^. Previous experiments have focused primarily on responses to rule deviants or “global deviant” responses, with the expectation that violation of a pattern should evoke a stronger response. Our neuronal results suggest a more nuanced understanding: responses to new exemplars of a familiar rule were at least as common as responses to rule deviants, implying that active pattern recognition may be just as relevant as deviant detection in these regions. These results are consistent with previous work on generalization of abstract category structure in LPFC^49,50^, extended to a sequence context. As we work towards understanding the neural processes that underpin these behaviors, establishing a functionally homologous primate model of abstract sequence tracking will be particularly valuable, opening the path for causal manipulation, behaviorally rich task design, and systematic investigation of the networks that enable complex cognition and behavior across species.

## Acknowledgements

We thank Matthew Maestri for his assistance with animal training and human and animal data collection throughout the course of this research. We thank Dr. David Sheinberg for advice and technical support on the task and software. We thank Dr. Michael Worden, Lynn Fanella, Dr. Edward Walsh, Fabienne McEleney, and other members of Brown University’s MRI Facilities staff for their support and guidance throughout this project.

## Contributions

Conceptualization: KEC, NYR, DB, THM, TMD. Monkey fMRI data acquisition: NYR, AA. Human fMRI data acquisition: MVRN, HH, THM, DB, NYR. Electrophysiology data acquisition: KEC. Data analysis: KEC, NYR, BS, MVRN, HH. Manuscript Preparation: KEC, TMD, BS, MVRN. Correspondence and material requests should be addressed to Dr. Theresa Desrochers.

## Methods

### Animal subjects

All procedures followed the NIH Guide for Care and Use of Laboratory Animals and were approved by the Institutional Animal Care and Use Committee (IACUC) at Brown University.

Three adult male rhesus macaques (Macaca mulatta) were used in fMRI experiments (monkeys B, J, and W; ages 6-12 years old during data collection; weights 7.4-16.5kg). Before task training, each monkey was surgically implanted with an MR-compatible head-restraining device (PEEK, Applied Prototype). After fMRI data collection, two of the macaques were implanted with custom PEEK chambers for electrophysiology recording (Rogue Research). Craniotomies were performed during the same surgery as chamber implantation. The final position and angle of the chamber were confirmed using post-surgery structural MRI scans with the saline-filled chamber (monkey W: left hemisphere, centered at - 15.5mm ML and +33.5mm AP, with the top of the chamber angled 33.5° anterior and 10° medial; monkey B: right hemisphere, centered at +16.0mm ML and +36.5mm AP, with the top angled 22.8° anterior and 10° medial). All surgical procedures were performed under general anesthesia with analgesics delivered post-operatively. A minimum of 8 weeks recovery time was given after head-holder or chamber implantation surgery before beginning training or data collection.

### Human participants

Participants were recruited via email advertising, flyers, and word of mouth. Inclusion criteria were as follows: age 18-35 years old, right-handed, vision normal or corrected-to-normal, no psychiatric or medical diagnoses, and no contraindications for fMRI. All participants gave informed, written consent, and study procedures were approved by the Brown University Institutional Review Board. Participants completed an initial in-person training visit to be familiarized with the abstract sequence viewing task used in the study. Training consisted of two task runs with eye tracking, and participants received $10 per hour for completing the session. After performing the task, subjects were asked to identify the sequence rule used in the training runs (“OneDiff” or “AllSame”, see “*Task Structure”*), and those who could do so accurately were invited to participate in the fMRI session. The fMRI portion consisted of one visit of 5-6 task runs with eye tracking (see “*Task Structure*”). At the end of the scan, participants completed a questionnaire that asked them to identify any rule(s) they noticed and stimuli they observed. Participants were paid $20 per hour for the fMRI scan visit. Initially, 39 participants were recruited to complete task training and the scanning experiment. A total of 10 participants were excluded due to excessive motion (translation >3mm displacement and rotation >2°) and one was excluded due to poor fixation (fixating <80% of the time per run on average). After excluding participants, our final sample size included 28 participants (mean age 22.7 (+/-5.1 [SD]); 16 males [12 f]). This sample size exceeds the number determined by a power analysis needed to achieve 80% power with an effect size of 0.5 (Cohen’s d) in this study (n = 26).

### Task structure

All experiments measured responses in the lateral prefrontal cortex during variants of a no-report abstract sequence viewing task, described in detail below. In this task, subjects fixated on the center of a gray screen as fractal images were presented sequentially. The general design of this paradigm was based on a similar auditory sequence task^1^.

#### Stimuli

Visual stimuli in all tasks were displayed using a custom software system based on OpenGL, developed by Dr. David Sheinberg at Brown University. The experimental task was controlled with custom Tcl code on a QNX real-time operating system using a state machine with a C backend. Eye position was monitored using video eye tracking (Eyelink 1000, SR Research) and was calibrated before data collection each day (9-point calibration with double quadratic regression). Stimuli in fMRI experiments were displayed on a 24-inch BOLDscreen flat-panel display (Cambridge Systems). Monkeys were scanned in the sphinx position and viewed the screen directly. Humans were scanned in a supine position, with the display and eye tracking reflected through a mirror box mounted on the head coil. Stimuli in electrophysiology experiments were displayed on a 32-inch LCD monitor (LG; 144Hz refresh rate) centered at eye level 100cm in front of the animal while the monkey sat upright in a custom chair.

Each stimulus consisted of a fractal (∼8° visual angle) with varying colors and features. Fractals were generated using MATLAB for each scanning session using custom scripts based on stimuli from Kim and Hikosaka^2^, following the instructions outlined in Miyashita et al.^3^. Each stimulus was displayed for 0.1, 0.2, or 0.3 s, depending on the timing template for the current sequence (see “*Sequence types”*, Supplementary Fig.1A). All stimuli were presented on a gray background. A small fixation spot (0.3° square) was present on the center of the screen throughout the task and was superimposed on the fractal images when they appeared. To provide behavioral feedback, the fixation spot was yellow while the subject maintained fixation within a small window (typically 3° of visual angle) and turned red when the subject was not fixating.

#### Sequence types

FMRI experiments used five types of sequences: habituation sequences and four types of deviant sequence. Electrophysiology experiments used habituation sequences and two types of deviant. Habituation and deviant sequences in all tasks used separate pools of images. New, luminance-matched fractal sets were generated for each day of data collection, and separate fractals were used for fMRI and electrophysiology experiments. Behavioral training for fMRI experiments was conducted using a standard set of scrambled object images, and training for the electrophysiology experiments was conducted using a separate set of fractal images. We refer to the habituation images as [A, B, C, D] and the deviant images as [E, F, G] (Fig.1D). Each sequence can be categorized based on the abstract pattern it follows across stimuli, which we refer to as the “sequence rule”. Four item-sequences in this task each followed one of two patterns: “four repetitions of the same image” (“AllSame”) or “three of the same image followed by one different” (“OneDiff”).

#### Habituation sequences

Habituation sequences were four-item sequences that followed a consistent pattern throughout the task run. In AllSame runs, the rule was “four repetitions of the same image” (e.g. AAAA, CCCC). In OneDiff runs, the rule was “three of the same, then one different” (e.g. AAAB, DDDC). Habituation sequences had three possible durations: short (Hs, 1.1s), medium (Hm, 1.7s), or long (Hl, 2.3s). For each total duration, there were two possible stimulus timings: one with a shorter stimulus presentation and longer inter-stimulus interval (ISI), and one with a longer stimulus duration and shorter ISI. This arrangement gave a total of six possible timings: short1 (stimulus duration: 0.1s, ISI: 0.2333s), short2 (stimulus duration: 0.2s, ISI: 0.1s), medium1 (stimulus duration: 0.1s, ISI: 0.4333s), medium2 (stimulus duration: 0.3s, ISI: 0.1667s), long1 (stimulus duration: 0.2s, ISI: 0.5s), and long2 (stimulus duration: 0.3s, ISI: 0.3667s) (Fig.S1A). Habituation sequences were used in all experiments.

#### New Images with the Same Rule (NI)

NI sequences followed the same pattern as habituation sequences but used fractals drawn from the deviant image pool. For example, in AllSame runs, GGGG would be an NI deviant; in OneDiff runs, GGGF would be an NI deviant. Each fractal was presented for 0.2s with an ISI of 0.3s, for a total duration of 1.7s (matching the total duration of medium habituation sequences). NI sequences were used in all experiments.

#### Rule Deviants (RD)

Rule Deviants were four-item sequences that did not match the habituation sequence pattern and instead used the other sequence rule. For example, in AllSame runs, the habituation rule was “all four of the same image” (e.g. AAAA), but Rule Deviants would be “three of the same, then one different” (e.g. GGGF). In OneDiff runs, the opposite was true, and Rule Deviants followed the pattern “all four of the same” (e.g. GGGG). Rule Deviants used the same timing as NI sequences: 0.2s image duration, 0.3s ISI, and 1.7s total sequence duration. RD sequences were used in all experiments.

#### Number Deviants (ND; fMRI only)

Number deviants had either two images or six images instead of four, but otherwise followed the standard rule for the run. Thus, in AllSame runs, Number Deviants would instead follow the rule “two of the same image” (e.g. FF) or “six of the same image” (e.g. FFFFFF). In OneDiff runs, Number Deviants would either be “one image followed by one different image” (e.g. FG) or “five of the same followed by one different image” (e.g. FFFFFG). Images were presented for 0.2s each in both types of Number Deviant. Two-item NDs had an ISI of 0.7s, for a total duration of 1.1s (matching the total duration of short habituation sequences). Six-item NDs had an ISI of 0.3667s, for a total duration of 2.3s (matching the total duration of long habituation sequences). NDs were only included in human and monkey fMRI variants of the task, not in electrophysiology experiments.

#### Double Deviants (DD; fMRI only)

Double deviants differed from habituation sequences in both sequence length and the sequence pattern. As with Number Deviants, they could be either 2-item or 6-item, but they followed the opposite sequence pattern rule. Thus if the habituation pattern was “all four the same,” Double Deviants were either “one image followed by one different image” (e.g. FG) or “five of the same followed by one different image” (e.g. FFFFFG). The timing for Double Deviants was the same as Number Deviants. Like Number Deviants, Double Deviants were included only in the fMRI variants of the task, and not for primate electrophysiology. We note that Double Deviants were included to balance the two-by-two design and to parallel a previous auditory task^1^. However, Double Deviants were not used in subsequent analyses because of the inability to fully differentiate the effects of rule and number.

#### Run structure

Each day of data collection consisted of several task runs (monkey fMRI: 4-8 runs/day; human fMRI: 5-6 runs/subject; electrophysiology: 4-16 runs/day). Each run was defined by one of two standard sequence patterns: AllSame (four repeated images; e.g. AAAA, BBBB); or OneDiff (three of the same image followed by one different; e.g. AAAB, DDDC). The standard pattern was counterbalanced across runs to maintain an approximately equal number of runs for each rule on each day. The same run structure was used for behavioral training prior to data collection.

Each run lasted approximately 10 minutes, and consisted of five blocks of thirty trials each, interleaved with 14s fixation miniblocks with no fractal stimuli. We defined one trial as a complete sequence (2, 4, or 6 fractal stimuli, depending on the sequence type). The first block (“habituation”) consisted of 30 habituation sequences. In each subsequent block (blocks 2-5), 80% of trials had habituation sequences (24 sequences), but 20% of trials used one of the four types of deviant sequences (6 sequences). Each block only contained one type of deviant sequence - i.e., a “Rule Deviant” block consisted of 80% habituation sequences and 20% Rule Deviants, but no other sequence types.

In human and monkey fMRI experiments, each of the four deviant types were used for one of the four blocks after habituation, with the order counterbalanced across runs. Number and Double Deviant blocks each contained three short and three long deviants. Electrophysiology experiments used only NI and RD blocks after habituation. In the majority of electrophysiology runs, NI and RD blocks alternated after the initial habituation block, with the first deviant block type determined pseudorandomly. However, due to code error, OneDiff runs from monkey W included three RD blocks and only one NI block. In these runs, the NI block was always either the fourth or the fifth block of the run.

Within a block, the order of sequences was constrained in the following ways: 1) the first fractal in one sequence could not match the final fractal of the previous sequence; 2) the first six sequences in any block had to be habituation sequences; 3) two deviant sequences could not be presented back-to-back; 4) for ND and DD blocks, an equal number of short (2-item) and long (6-item) deviants were used.

The intersequence interval was jittered to decorrelate responses across different sequence types (mean 2s, 0.25-8s). For consistency, the same distribution of intersequence intervals was used across all task variants (human fMRI, monkey fMRI, and monkey electrophysiology).

#### Reward schedule

In both monkey fMRI and electrophysiology experiments, monkeys received liquid rewards for their engagement in the task. The timing of rewards was not linked to stimulus presentation or any other sequence events and depended exclusively on the monkey’s fixation. Rewards were delivered on a graduated schedule, such that the frequency of reward increased as the monkey maintained fixation for longer periods. To count as “fixation,” the monkey had to maintain gaze within a small window (typically 3° of visual angle) around the fixation point. The first reward was given after 4 s of continuous fixation. After two consecutive rewards, the fixation interval required for reward decreased by a small amount (0.25 s for electrophysiology experiments, 0.05 s for fMRI). The minimum inter-reward interval that a monkey could reach was 0.5 s, though this was not typically achieved. In fMRI runs, 0.32 s allowance was provided for blinks. If the monkey’s eyes left the fixation window and returned within that time allowance, it would not trigger a fixation break. For the electrophysiology experiment, a blink allowance of 0.64 s was provided for monkey B, and no blink allowance was provided for monkey W. If fixation was broken for longer than the allowance, the reward schedule would reset to the maximum 4 s fixation duration required to obtain reward.

In human fMRI experiments, no performance-based reward was given. To provide a signal that matched timing of reward events, the central fixation spot flashed green if the human participant maintained fixation within the fixation window, with timing that matched the reward schedule in macaque fMRI variant (4 s initial duration, decreasing by 0.05 s after every two successful intervals). No blink allowance was provided for human participants. Participants were informed that the green flash reflected good performance on the task, but the color change signal was not correlated with monetary rewards.

### MRI Data Acquisition

#### Monkey fMRI data acquisition

Methods for this experiment were described in detail in Yusif Rodriguez et al.^4^ and are summarized here. Monkeys were habituated to all scanning procedures for several weeks prior to data collection. During fMRI experiments, monkeys sat in the “sphinx” position in an MR-safe primate chair (Applied Prototype or custom-made by Brown University) and wore earplugs (Mack’s Soft Moldable Silicone Putty Ear Plugs, “kid’s” size). A plastic headpost (PEEK, Applied Prototype) was used to restrain their head during data acquisition. Approximately 30–60 min prior to each scanning session, monkeys were intravenously injected with a contrast agent: monocrystalline iron oxide nanoparticle (MION; either Feraheme (ferumoxytol), AMAG Pharmaceuticals, 30 mg per ml, or BioPAL Molday ION, BioPhysics Assay Laboratory, 30 mg per ml). MION was injected into the saphenous vein below the knee (7 mg/kg) and then flushed with approximately double the volume of sterile saline. No additional MION was added during scanning.

A Siemens 3T PRISMA MRI system with a custom six-channel surface coil (ScanMed) at the Brown University MRI Research Facility was used for whole-brain imaging. Anatomical scans consisted of a T1-MPRAGE (repetition time [TR], 2,700 ms; echo time [TE], 3.16 ms; flip angle, 9°; 208 sagittal slices; 0.5 × 0.5 × 0.5 mm), a T2 anatomical (TR, 3,200 ms; TE, 410 ms; variable flip angle; 192 interleaved transversal slices; 0.4 × 0.4 × 0.4 mm), and an additional high-resolution T2 anatomical (TR, 8,020 ms; TE, 44 ms; flip angle, 122°; 30 interleaved transversal slices; 0.4 × 0.4 × 1.2 mm). Functional images were acquired using a fat-saturated gradient-echoplanar sequence (TR, 1.8 s; TE, 15 ms; flip angle, 80°; 40 interleaved axial slices; field of view [FOV] read, 92mm; FOV phase, 100%; slice thickness, 1.1mm; matrix, 84 x 84; 1.1 × 1.1 × 1.1 mm).

#### Human fMRI data acquisition

Whole-brain imaging was conducted at the same facility using the same scanner as for monkeys but with a 32-channel head coil. Functional data for all participants were acquired using a gradient-echoplanar imaging pulse sequence (TR, 2.0 s; TE, 28 ms; flip angle 90°; 38 interleaved axial slices; FOV read, 192mm; FOV phase, 100%; slice thickness, 3.0mm; matrix, 64 x 64; 3.0 x 3.0 x 3.0 mm). Anatomical scans included a T1-MPRAGE (TR, 1900 ms; TE, 3.02 ms; flip angle, 9.0°; 160 sagittal slices; 1.0 x 1.0 x 1.0 mm) and a T1 in-plane scan (TR, 350 ms; TE 2.5 ms; flip angle, 70°; 38 transversal slices; 1.5 x 1.5 x 3.0 mm).

### FMRI Analysis

Analyses for both human and monkey fMRI experiments were performed in MATLAB 2025a using Statistical Parametric Mapping (SPM12) (http://www.fil.Ion.ucl.ac.uk/spm).

#### Monkey fMRI analysis

Preprocessing and data inclusion criteria were similar to previous analyses of this dataset^4,5^. Preprocessing was performed prior to data analysis, and consisted of reorienting (to ensure proper assignment of the *x*, *y*, *z* planes), motion correction (realignment), normalization, and spatial smoothing (3 mm isotropic Gaussian kernel separately for gray matter and white matter). All steps were performed on individual runs separately. The T1-MPRAGE anatomical image was skull stripped using FSL BET brain extraction tool (http://www.fmrib.ox.ac.uk/fsl/) to facilitate normalization. For most analyses, images were normalized to the 112RM-SL macaque atlas^60^. Analyses for monkeys W and B were repeated in each monkey’s native space to facilitate fMRI-guided electrophysiology and comparisons across different recording sites (see “*Target Selection*” and “*Analysis of recording site effects”*). Runs were only included for analysis if the monkey was fixating for >80% of the task and a sufficient number of acquisition volumes passed data quality checks. We used the ART toolbox (Artifact Detection Tools, https://www.nitrc.org/projects/artifact_detect) to detect outlier volumes (standard global mean; global signal detection outlier detection threshold, 4.5; motion threshold, 1.1 mm; scan to scan motion and global signal change for outlier detection). Any run with >12% of volumes detected as outliers was excluded from analysis. Runs with image quality too poor to preprocess or analyze because of artifacts or banding were also excluded. After applying these criteria, a total of 232 runs (71 sessions in total across animals) were included for analysis from three monkeys: monkey W: 97 runs (28 sessions), monkey J: 65 runs (27 sessions), and monkey B: 70 runs (16 sessions).

#### Monkey fMRI model

Analyses were performed using pseudo-subject bins consisting of approximately 10 runs/bin. Each bin contained data from only one monkey. Runs were pseudo-randomly assigned to bins to balance the proportion of different run patterns (OneDiff vs. AllSame) and the distribution of runs from earlier and later scanning sessions. Bin assignments were slightly changed compared to previous analyses^4^ to ensure that no bins had >60% of data from one run pattern.

Within-subject statistical models were constructed under the assumptions of the general linear model (GLM) in SPM12 for each pseudo-subject bin. Condition regressors were all convolved with a γ function (shape parameter = 1.55, scale parameter = 0.022727) to model the MION hemodynamic response function^61^. The first six sequences in each run and reward times were included as nuisance conditions. Additional nuisance regressors were included for the six motion estimate parameters (translation and rotation), outlier volumes, and image variability (SD of within run image movement variability, calculated using the ART toolbox). Outlier volumes were determined using the ART toolbox (standard global mean; global signal detection outlier detection threshold = 4.5; motion threshold = 1.1 mm; scan to scan motion and global signal change for outlier detection), and the model included one additional regressor with a “1” at each volume to be “scrubbed.”

Regressors were estimated using a bin-specific fixed-effects model. Whole-brain estimates of bin-specific effects were entered into second-level analyses that treated bin as a random effect. Significance was assessed using one-sample one-sided t-tests (contrast value vs. zero, p < 0.005), corrected for multiple comparisons at a cluster level to yield family-wise error (FWE) error correction (p < 0.05). To assess the univariate effects of deviant sequences, we constructed a GLM using instantaneous stimulus onset regressors for the first item in each sequence with the following nine condition regressors for different sequence types: short, medium, and long habituation sequence timing templates; new images with the same rule; rule deviants; two-item and six-item number deviants; and two-item and six-item double deviants.

#### Human fMRI analysis

Preprocessing steps included slice time correction, motion correction (realignment, B-spline interpolation), normalization (Montreal Neurological Institute [MNI] template with affine regularization), and spatial smoothing (8-mm full-width at half-maximum Gaussian kernel and resampled using trilinear interpolation), to largely parallel preprocessing of monkey data.

#### Human explicit masking

Subject-level brain masks were generated using FastSurfer^6^, a deep learning-based skull-stripping and segmentation pipeline that was applied to each participant’s T1-weighted anatomical image. The mask was then warped into normalized space using each participant’s deformation field and boundary restricted to the acquired functional field of view. The convergence mask allowed us to (a) exclude any anatomically-labeled voxels not collected during acquisition, and (b) utilize it as an explicit mask during first-level model estimation. Since masking was handled entirely by the explicit convergence mask, SPM12’s default relative intensity threshold (i.e., masking threshold applied as a proportion of the global mean signal) was disabled (mthresh = 0). This approach was adopted after observing that the conventional threshold approach (i.e., excluding voxels below 80% of the global mean) removed low-intensity yet valid cortical tissue in regions of interest, such as the orbitofrontal and rostral prefrontal cortex, while a fully permissive threshold (0%) allowed non-brain voxels into the estimation, producing a degenerate noise model. Restricting the explicit mask to anatomically-defined and functionally-sampled tissue avoided both problems.

#### Human fMRI model

GLMs were constructed using SPM12 and custom scripts in MATLAB. Onset (zero duration) regressors at the first stimulus in each sequence were convolved with the canonical hemodynamic response function (HRF) and the first time derivative of the HRF. Separate regressors were included for the following nine conditions: short, medium, and long habituation sequence timing templates; new images with the same rule; rule deviants; two-item and six-item number deviants; and two-item and six-item double deviants. Nuisance regressors were included to account for variance due to motion (translation and rotation), the first six trials of every block, and the timing of the green color change of fixation spot.

Beta values related to regressors were estimated using a subject-specific fixed-effects model. Whole brain contrasts estimated subject-specific effects, and these estimates were entered into a second-level analysis with subject treated as a random effect. Whole-brain group voxel-wise effects were thresholded at p < 0.005 (uncorrected) to form clusters, and corrected for multiple comparisons using cluster-extent thresholds to yield FWE correction. Clusters were considered significant at p(FWE) < 0.05. Group level contrasts were rendered on MRIcron (https://www.nitrc.org/projects/mricron) and MRIcroGL (https://www.nitrc.org/projects/mricrogl/). The same RD>NI onsets model was used for the human and monkey GLMs (see “*Monkey fMRI model”*).

#### Inter-Species fMRI correlation

Functional interspecies correlations (ISC) were assessed similarly to Caspari et al.^7^ Voxelwise beta parameters quantifying the impact of experimental conditions on brain activity were extracted from the GLMs described above for short habituation (Hs), medium habituation (Hm), long habituation (Hl), new images with the same rule (NI), 2-item number deviant (ND2), 6-item number deviant (ND6), and rule deviant (RD) sequences for each run from each human subject and monkey pseudo-subject bin. To account for beta parameter variability across runs, within-subject mean beta parameters were calculated for each condition and converted to *t*-values by dividing mean beta parameters by subject-specific beta standard errors for each condition (i.e. using the contrast of the condition over baseline). Group mean *t*-values were calculated by averaging *t*-values across subjects.

Arrays with *t*-values for each included experimental condition were generated for each brain voxel. Region of interest (ROI) arrays were generated by averaging condition specific *t*-values across all contained voxels. ROI-to-brain correlations compared source species group-average ROI *t*-value arrays with voxelwise *t*-value arrays from each target species subject. Pearson’s *r* was calculated for the correlation at each voxel, followed by Fisher’s r-to-z transformation, generating a z-correlation map for each target species subject. Group-level significance of correlations was determined by entering each target species subject z-correlation map into a random-effects analysis thresholded a p < 0.005, followed by family-wise error (FWE) rate cluster correction to threshold at p-FWE < 0.05.

### Extracellular Recording

Extracellular recordings were collected using a Grapevine Neural Interface Processor and Trellis Neural Acquisition Software Suite (Ripple Neuro). The spike datastream was amplified, bandpass filtered (high-pass cutoff: 250 Hz; low-pass cutoff: 75 kHz) and sampled at 30 kHz. Spike events were saved any time the signal crossed a user-defined voltage threshold (typically ∼25-30uV). Recordings were synchronized with the behavioral task using a TTL pulse sent from the behavior control system at the beginning and end of each task run. Data from monkey B were collected using 32-channel linear electrodes (Plexon V-probes, 32-channels, 50µm spacing between sites). Data from monkey W were collected using both single tungsten electrodes (FHC) and multisite linear electrodes (Plexon V-Probes, 16-32 channels, 75-150µm spacing between sites), depending on the recording day. Electrode position was set using an XY-positioner (NAN instruments; monkeys B and W) or 1mm-spaced grid (Rogue Research; monkey W). Guide tubes were used to penetrate the dura before recording each day, and electrodes were lowered using a hydraulic microdrive (Kopf) or electronic motor (NAN Instruments).

On recording days, electrodes were lowered along a planned trajectory to the approximate target depth, with the specific recording depth chosen based on the presence of clear neuronal signals. V-probe recordings were raised ∼100µm after the initial lowering and allowed to settle for ∼30 minutes before data collection to improve the stability of signals over time. For V-probe recordings and a subset of tungsten electrode recordings, multiple runs were collected at the same recording depth (minimum of 4 runs per depth). For the remainder of tungsten electrode recordings, electrode depth was changed between runs until the signals from previous cells were absent and new units could be clearly detected (≥50µm difference between runs). Spike sorting was performed offline using Plexon Offline Sorter.

#### Target selection

Recording targets were selected based on fMRI data from each monkey. Rather than normalizing to the atlas, images for each monkey were aligned to their native anatomical space. Models were run separately for monkey W and monkey B, using only the data that had been collected for that monkey. Contrasts were created for RD>NI Onsets for each individual monkey, using a liberal threshold (p < 0.05 uncorrected) for targeting purposes. Guided by these contrasts, recording trajectories were planned using Brainsight neuronavigation software (Rogue Research) to sample neurons in and around fMRI-defined regions of interest (Fig.3A, Supplementary Fig.3). After data collection, we determined the specific location of each recording site based on the penetration site, depth from tissue surface, and the chamber position and angle.

#### Duplicate removal

To avoid double-counting, each pair of simultaneously recorded units was checked for duplicate spikes, defined as spikes with <8.3 µs difference in timestamp across channels. Units that shared >20% of spikes with another channel were considered cross-contaminated. The median waveform parameters for duplicate spikes were compared to non-duplicate spikes for each unit in the pair (parameters: peak-to-valley amplitude, valley full-width-at-half-maximum, and peak full-width-at-half-maximum). Duplicate spikes were assigned to the unit with more similar waveform parameters and removed from the other unit.

### Electrophysiology Analysis

Analyses were conducted in MATLAB. We analyzed responses in time windows 50-350 ms after the appearance (ImgOn) and disappearance (ImgOff) of each fractal. For units recorded across multiple task runs, firing rates were z-scored within each run and pooled across runs. Since the reward schedule was independent of sequence timing, rewards only occurred for a subset of stimuli and could happen at different times relative to stimulus onset and offset. To mitigate the effects of this variability, analyses excluded stimuli that overlapped with reward delivery. This constraint decreased variability related to movement noise and trial-to-trial differences reward timing. Including rewarded trials did not substantively change the core results.

#### Inclusion criteria

Units were included in analysis if they had stable waveforms with a mean firing rate of >1 sp/s in at least one run. Only runs with a mean firing rate >1 sp/s were analyzed. Trials were only included if the monkey maintained good fixation for the full duration of the sequence (>70% fixation for each of the four fractal images). If a monkey had bad fixation on >50% of sequences in a run, that run was excluded from analysis. For population analysis and decoding, units were only included if they included at least five good trials of each deviant type for both run patterns (OneDiff and AllSame) (Monkey W: 249 units; Monkey B: 444 units).

#### Single unit responses to task variables

Single-unit analyses focused on differences in firing rate across three primary task variables: ordinal position, sequence category (habituation vs. deviant), and deviant type (NI vs. RD). Responses to ordinal position and deviant status were identified using a two-way ANOVA (factors: ordinal position and deviant status, including interaction term). Responses to deviant type were identified using a separate one-way ANOVA on data from deviant trials only. We defined “task-related responses” as responses that showed significant differences in FR across one of these variables (p < 0.05). Additional one-way ANOVAs were run independently on data from habituation and deviant trials to identify response differences across specific fractal images. All tests were run separately for each neuron and time window.

#### Effect of image preference on deviant type responses

All units were included in the analysis, regardless of whether they showed significant selectivity for image or deviant type in individual ANOVAs. We defined each unit’s “preferred image” as the image that evoked the highest response, then computed the difference between RD and NI responses |*FR_RD_ - FR_NI_*| separately for the preferred image and non-preferred images. We compared the magnitude of the deviant-type effect on preferred vs. non-preferred images using a Wilcoxon sign-rank test. To avoid confounds, image preference was computed at sequence positions 1-3 and the magnitude effect was tested at position 4. A follow-up analysis confirmed that the magnitude effect was present at all four positions and was not limited to position 4.

#### Analysis of recording site effects

We used logistic regression to test whether the probability of finding task-related responses varied with distance from fMRI-defined regions of interest. For each monkey, we identified the largest clusters of voxels that showed fMRI responses to RD>NI in the recorded hemisphere (p < 0.05, uncorrected) and found the coordinates for the highest peak in that cluster (Supplementary Table 8). We then calculated the Euclidean distance of each recording site from the monkey-specific cluster peak. Using site distance as a predictor, we fit a GLM with a logit link function for each task variable: *logit(P(Y=1)) = β_0_+β_1_X,* where *X* was distance of each unit from the cluster peak and *Y* was a binary variable indicating whether the unit encoded the variable of interest. We fit the models separately for each monkey, and considered the effect significant only if the model overall showed a significantly better fit than a constant model (likelihood ratio test) and if the coefficient *β_1_*was significantly different from zero (Wald test).

#### Decoding

Decoding was performed using a pseudopopulation approach^8^ with a linear discriminant analysis classifier (LDA). The analysis used sliding time windows aligned to the sequence start, which covered the full duration of the sequence (time range: 50ms before image1 onset to 800ms after image4 offset; bin size: 200ms, step size: 10ms). As only deviant trials were included in analysis, stimulus onset and offset time were identical for all sequences. To obtain a cross-validated measure of decoder performance, data from all included units was split into random train and test sets (3:1 split), and separate groups of pseudotrials were constructed from each set. LDA was fit on pseudotrials drawn from the train set, then tested on pseudotrials from the held-out test set to give an accuracy estimate. For both the train and test set, pseudotrials were balanced across deviant type (RD vs. NI) and run pattern (OneDiff vs. AllSame), with 1000 pseudotrials sampled for each combination (OneDiff-RD, AllSame-RD, OneDiff-NI, and AllSame-NI). This process was repeated across ten iterations with different train/test splits, and the average cross-validated accuracy across these iterations was reported for each monkey. Time bins with significant above-chance decoding were identified using post-hoc one-sided t-tests on the pseudopopulation iterations with Benjamini-Hochberg FDR correction.

A contribution index for each unit was calculated for the time interval with the highest decoding accuracy using estimated Shapley values^9,10^. To estimate Shapley values, we generated 1000 randomly ordered permutations of the unit indices. For each permutation, units were added one at a time in the permuted order, and the pseudopopulation accuracy was computed for each added neuron. The overall contribution index of each neuron was defined as the change in decoding accuracy that occurred from adding that unit, averaged across all permutations. To test whether the decoding contribution was higher closer to the RD>NI cluster peak, we computed the Spearman correlation between contribution index and cluster distance and ran a permutation test to determine whether the correlation was significantly nonzero.

## Data and Code Availability

Code supporting the data analysis in this study will be made available on Github at the time of publication. Data that support the findings from the human participants in this study are available in the NIMH Data Archive (NDA) under Collection 4743, doi:10.15154/q92p-j128 and NDA Experiment 2590. Access is available to qualified researchers via an approved NDA Data Use Certification. Data that support the findings from nonhuman primates in this study are available from the corresponding author upon reasonable request.

## AI Use

Claude Code (Anthropic) was used in development of parts of the data analysis code. All code was manually reviewed and tested.

## Supplementary Figures

**Supplementary Figure 1.**
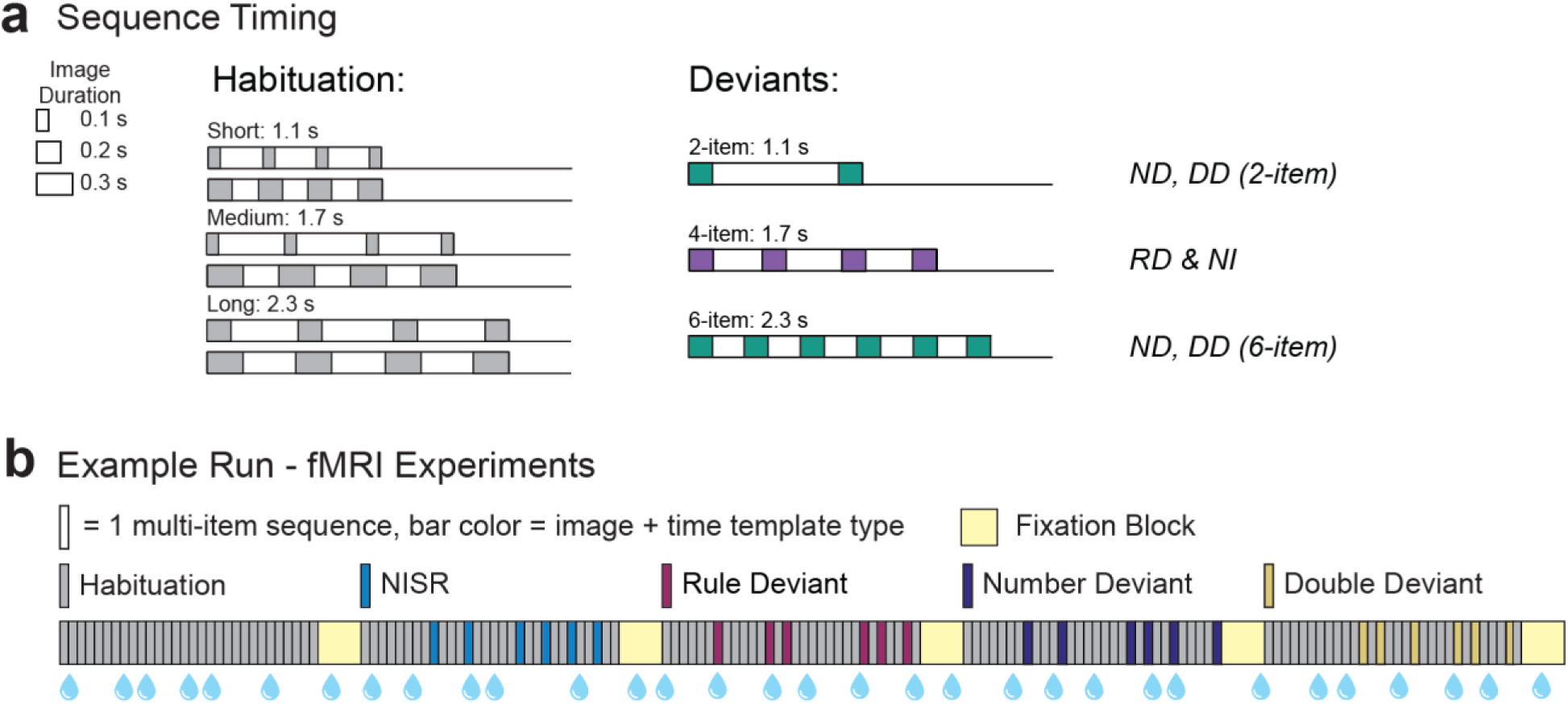
Details of the abstract sequence task. **(A)** Image and sequence timing for habituation and deviant sequences. Habituation sequences had six possible timings, which were interleaved in each block and used in all experiments. Deviant sequences were designed to match the total duration of either short, medium or long habituation sequences, with an image duration of 0.2s for each stimulus. FMRI experiments included all four deviant types. Electrophysiology experiments used only RD and NI deviants. RD = Rule Deviant, NI = New Images, ND = Number Deviant, DD = Double Deviant. **(B)** Schematic and example run from the monkey fMRI experiment. Each square denotes one multi-item sequence. The run consists of a block of 30 habituation sequences, followed by four blocks that show deviant sequences on 20% of trials. Each deviant type is present in one block. The order of deviant blocks was balanced across runs. Droplets indicate juice reward, which was delivered on a graduated schedule independent of sequence timing. The human fMRI task used identical run timing and structure, but did not include juice reward and instead used a green “reward signal” that matched the timing of juice delivery in the primate paradigm.

**Supplementary Figure 2.**
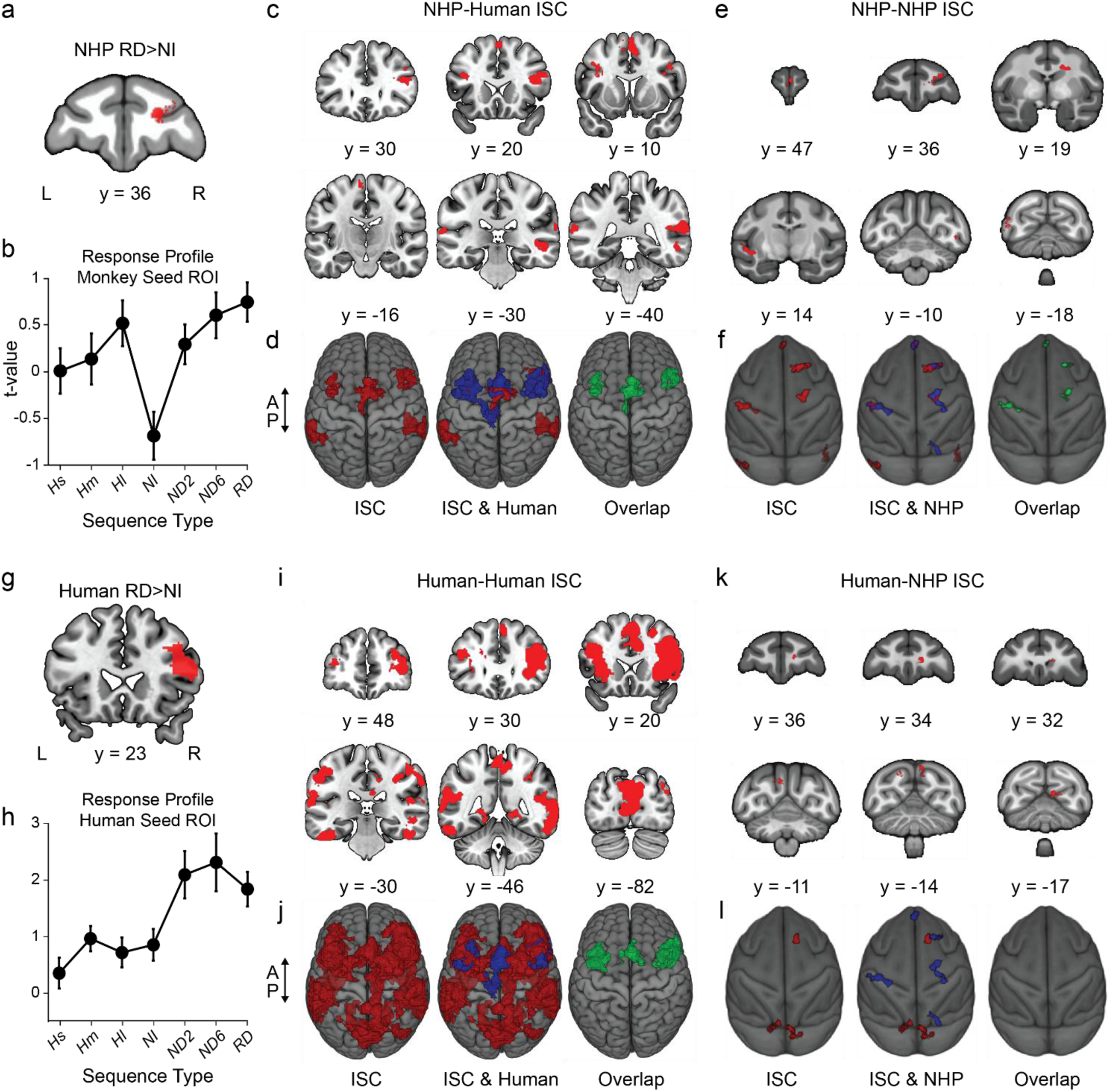
Interspecies and within-species fMRI correlation analyses. Panels A-D (enclosed in dashed gray border) are duplicated from Fig.2B-E for reference. **(A)** Monkey Rp46f subregion that responds more strongly to RD than NI sequences. **(B)** Activation profile in the Rp46f ISC seed region. **(C)** Human brain regions exhibiting responses across conditions significantly correlated with monkey Rp46f (FWEc < 0.05, height p < 0.005 unc., extent = 239). Error bars show ±SEM. **(D)** Rendered brains displaying monkey Rp46f-to-human ISC (red), monkey Rp46f-to-human ISC with the human RD>NI contrast from Fig. 2A (blue), and their overlap (green). (A-D) reproduced from Fig. 2B-E. **(E)** Monkey brain regions exhibiting responses across conditions significantly correlated with monkey Rp46f (FWEc < 0.05, height p < 0.005 unc., extent = 156); **(F)** rendered brains displaying monkey Rp46f-to-monkey ISC (red), monkey Rp46f-to-monkey ISC with the monkey RD>NI contrast (blue), and their overlap (green). **(G)** Human right LPFC subregion that responds more strongly to RD than NI sequences. **(H)** Activation profile in the right LPFC ISC seed region. **(I)** Human brain regions exhibiting responses across conditions significantly correlated with human right LPFC (FWEc < 0.05, height p < 0.005 unc., extent = 1237). Error bars show ±SEM. **(J)** Rendered brains displaying human right LPFC-to-human ISC (red), human right LPFC-to-human LPFC ISC with the human RD>NI contrast from Fig. 2A (blue), and their overlap (green). **(K)** Monkey brain regions exhibiting responses across conditions significantly correlated with human right LPFC (FWEc < 0.05, height p < 0.005 unc., extent = 184); **(L)** rendered brains displaying human right-LPFC-to-monkey ISC (red) and human right-LPFC-to-monkey ISC with the monkey RD>NI contrast (blue). These maps display no overlap.

**Supplementary Figure 3.**
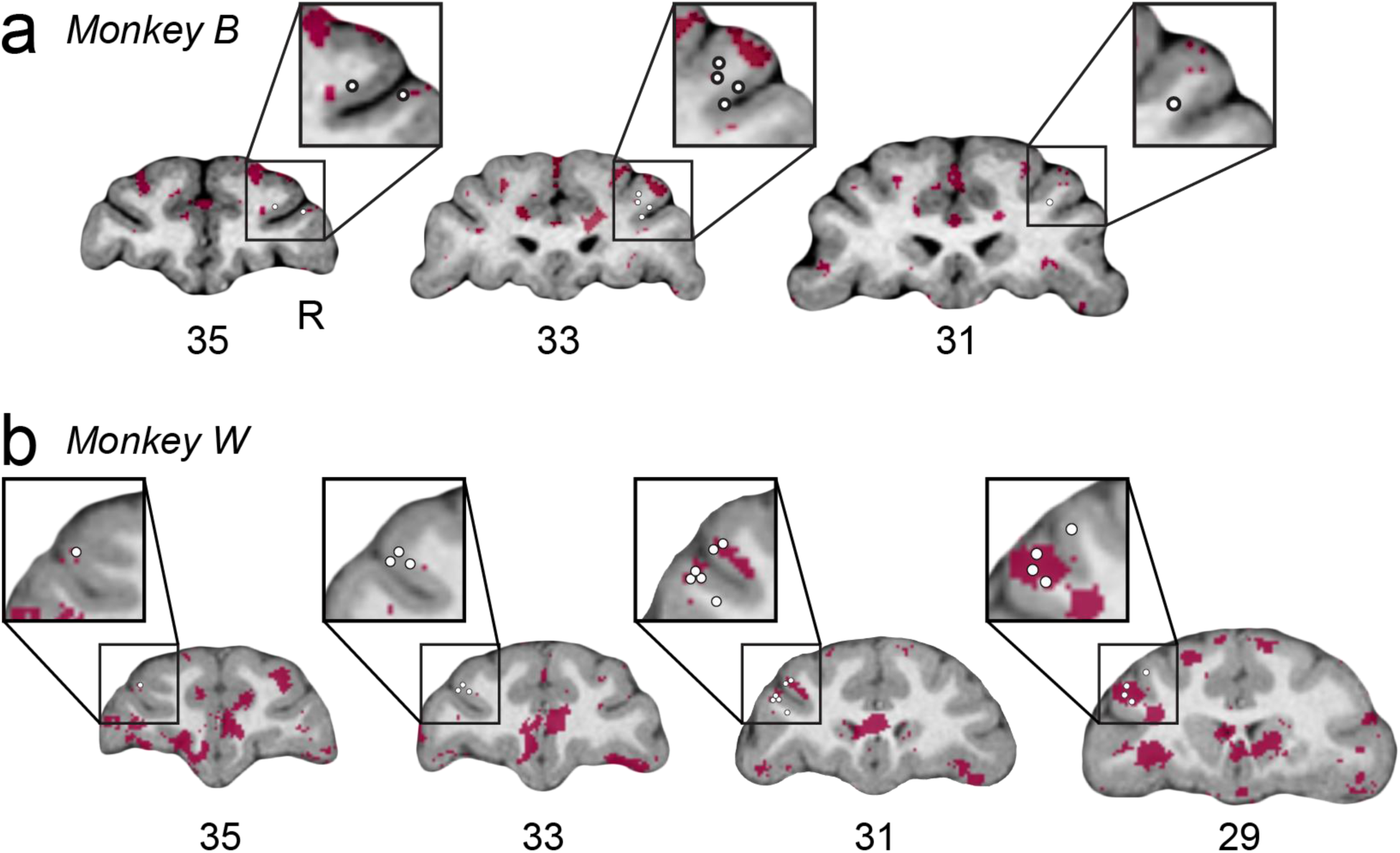
FMRI-guided recording sites from monkey B (A) and monkey W (B), displayed on a T1-weighted anatomical scan collected from the same monkey. White circles show the location of the deepest electrode site on a given recording day. Overlays show uncorrected fMRI contrasts for RD>NI onsets (maroon) (p<0.05, uncorrected). Slice numbers indicate the AP coordinate for each slice in the monkey’s native space.

**Supplementary Figure 4.**
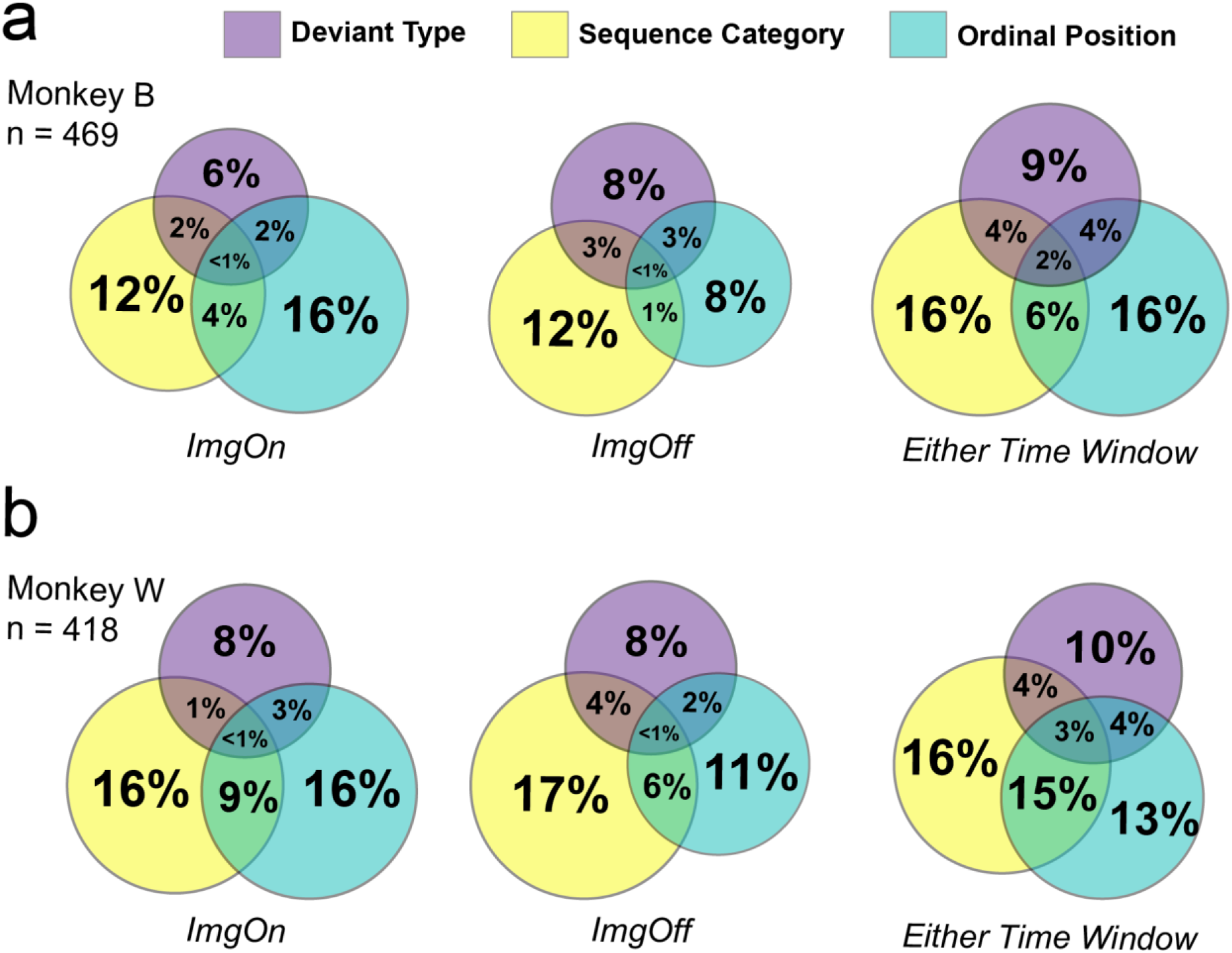
Prevalence and overlap of responses to deviant type, sequence category and ordinal position across LPFC neurons, separated by monkey and time window. Each Venn diagram shows the percentage of neurons that has a significant effect of one or more task variables (p<0.05; two-way ANOVA across sequence category and ordinal position; separate one-way ANOVA for deviant type).

**Supplementary Figure 5.**
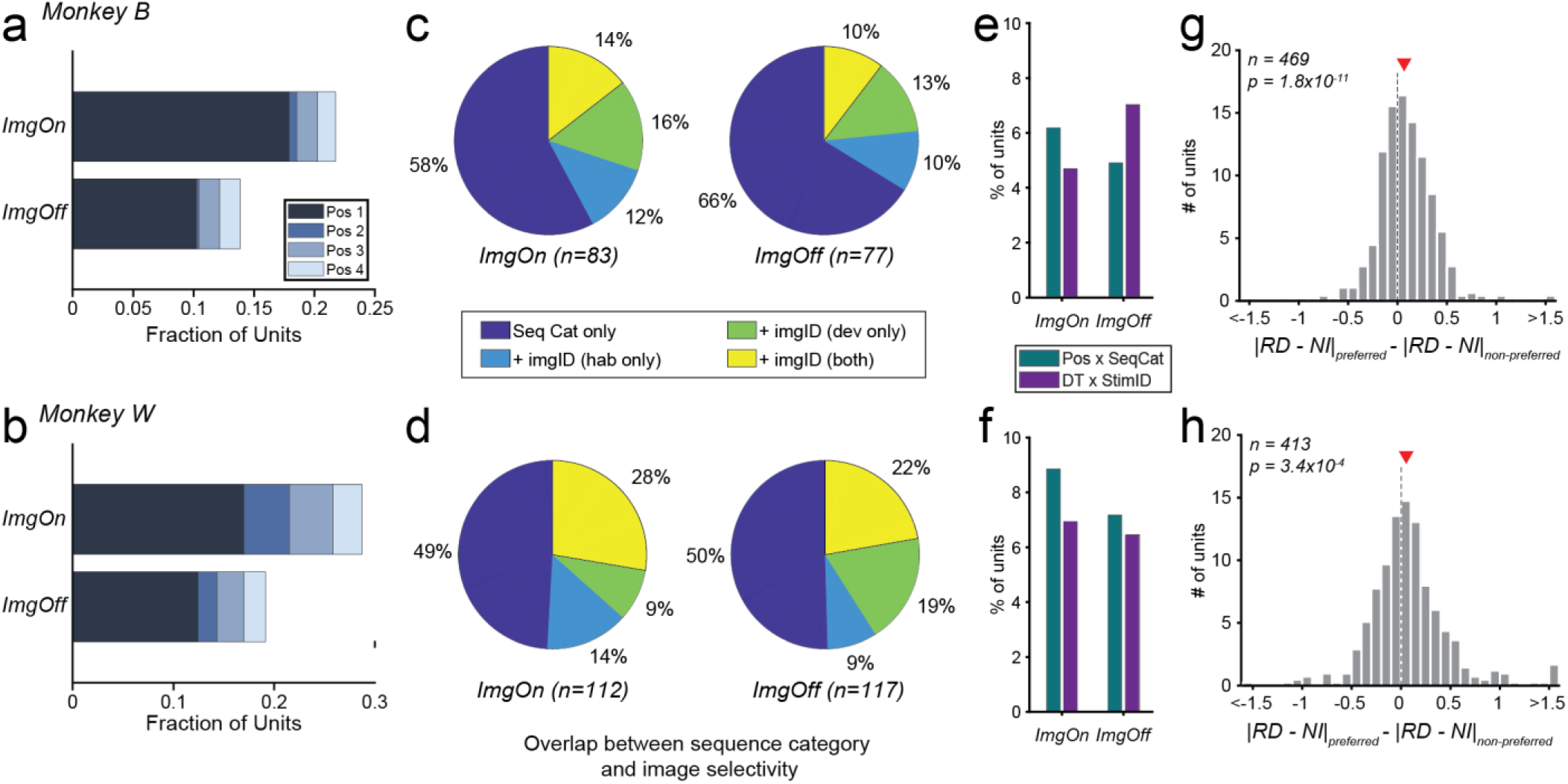
Details of task variable coding. Top: Monkey B. Bottom: Monkey W. **(A-B)** Fraction of units with a significant effect of sequence position, separated by the sequence position that they respond most strongly to (“preferred position”). More than half of position-selective units preferred position 1 in both monkeys and time windows. **(C-D)** Prevalence of significant within-pool image selectivity in units with a significant sequence category effect (habituation vs. deviant sequences). Image selectivity was tested with separate one-way ANOVAs in habituation and deviant sequences, since they used distinct pools of fractal stimuli. **(E-F)** Frequency of significant interaction effects for Ordinal Position x Sequence Category or Deviant Type (DT) x Image ID (p<0.05, separate two-way ANOVAs with interaction term). Interaction effects were uncommon in this dataset. **(G-H)** Effect of image preference on the magnitude of deviant type effects. Histograms show the difference in the absolute value of the deviant type effect |RD - NI| for each unit’s preferred stimulus minus its non-preferred stimuli (sign rank test). Red triangles mark the median value of the distribution; the dashed black line is zero. To minimize confounds, preferred stimuli were determined based on responses at positions 1-3, whereas RD and NI responses were taken from position 4. Units were included if there was at least one RD and one RD sequence with good behavior for each image in the deviant pool. Histograms show effects in the ImgOn time window. The effect was also significant in the ImgOff time window and when all four sequence positions were pooled (all p < 0.001, sign rank test).

**Supplementary Figure 6.**
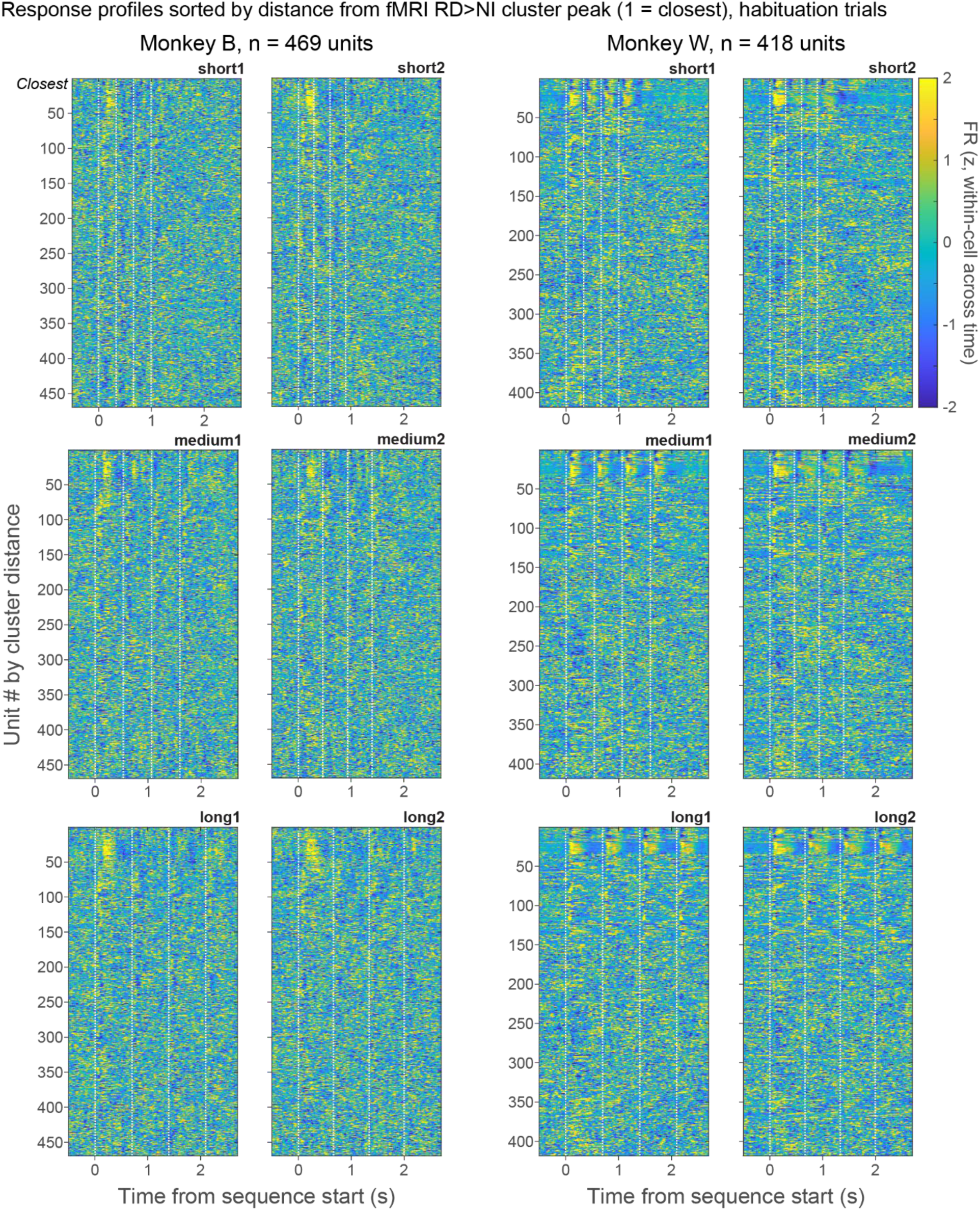
Neuronal responses for each of the six habituation sequence timings, sorted by distance from the RD>NI cluster peak. Responses were calculated as a sliding average with 50ms bins and 1ms steps.

**Supplementary Figure 7.**
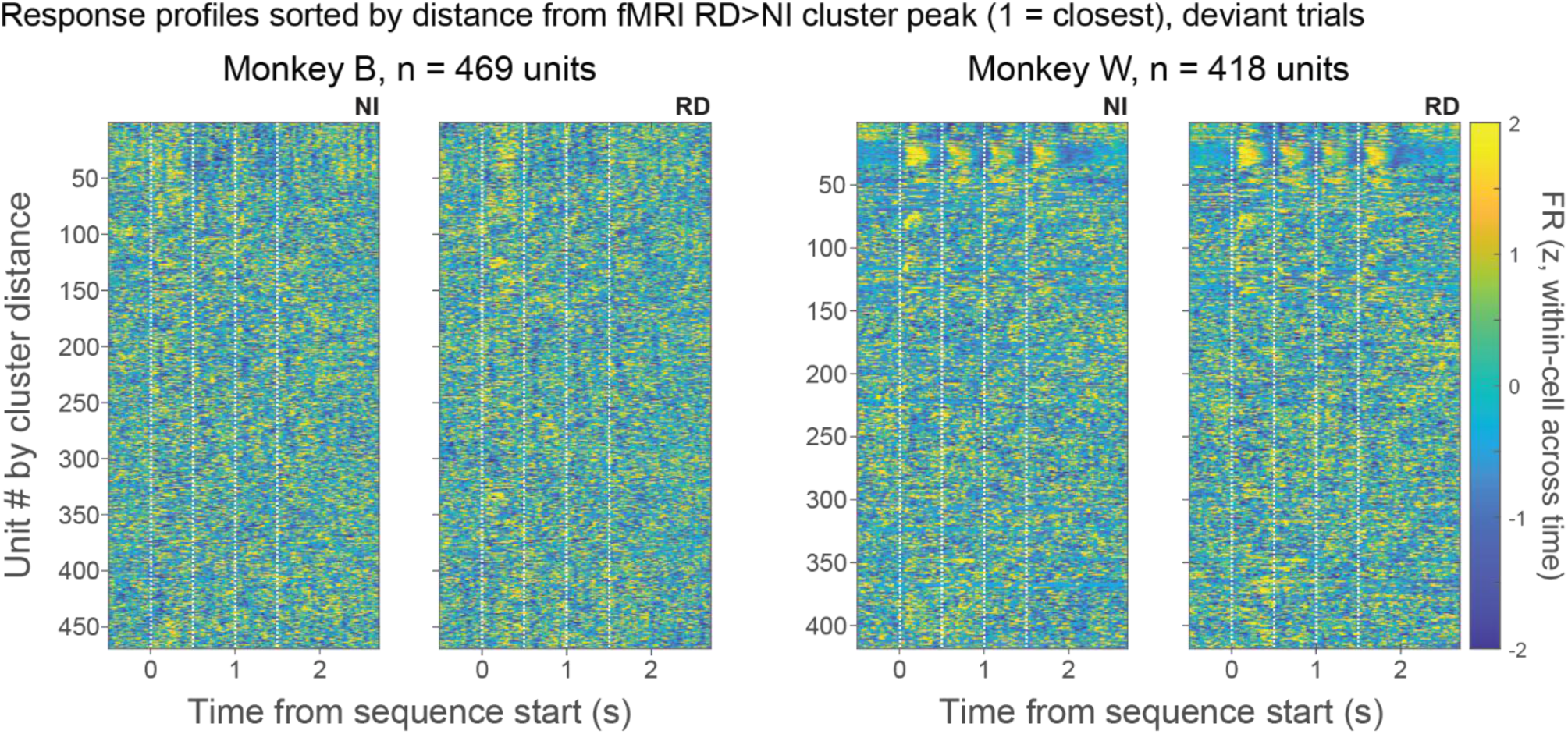
Neuronal responses for NI and RD sequences, sorted by distance from the RD>NI cluster peak. Responses were calculated as a sliding average with 50ms bins and 1ms steps.

**Supplementary Figure 8.**
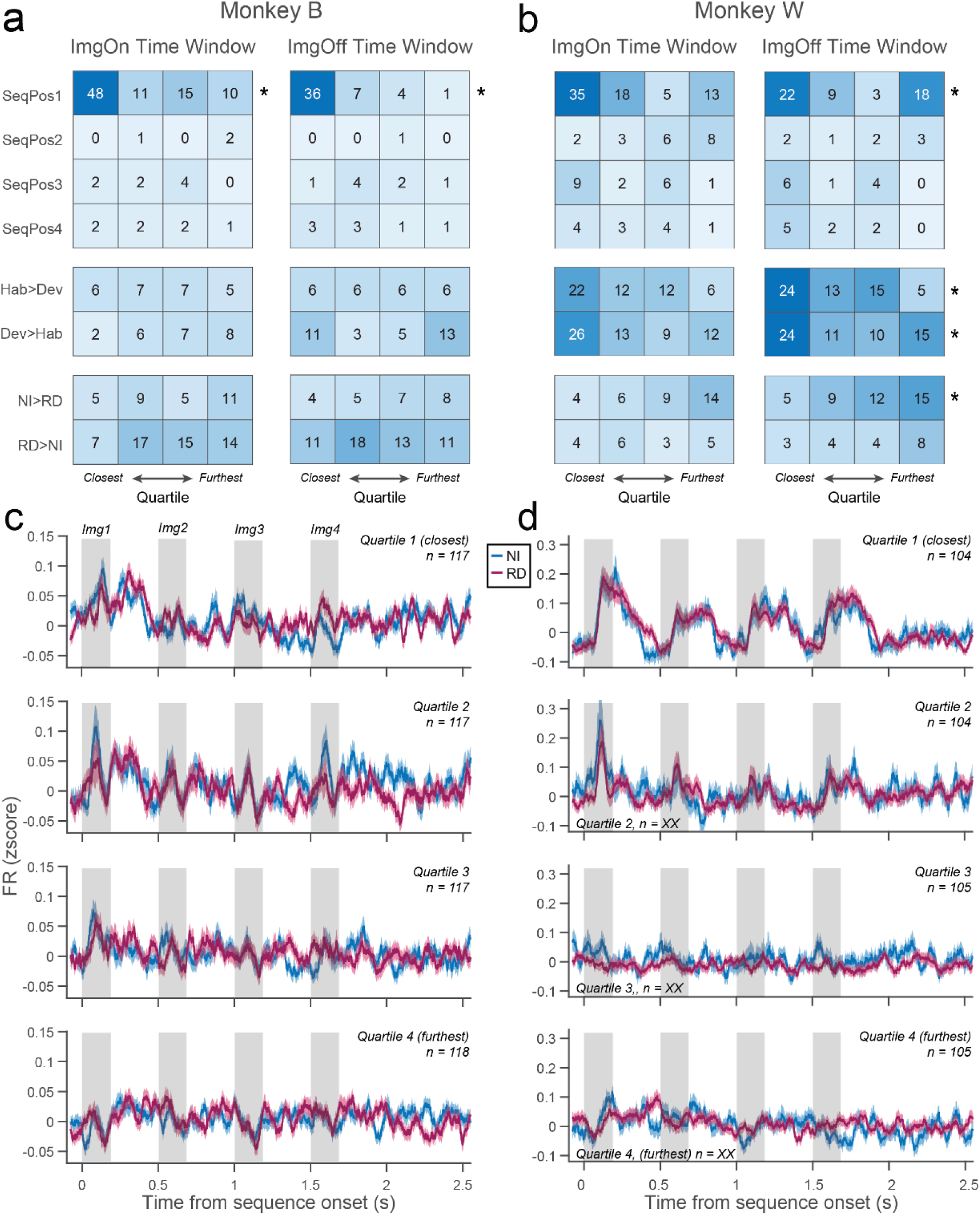
Neuronal response profiles change with distance from the RD>NI fMRI cluster peak. **(A-B):** Number of units that show significant effects for each task variable in each distance quartile, separated by monkey, preferred subtype, and time window. Preferred subtype is defined as the condition that evokes the highest response. Asterisks denote a significant effect of distance on the probability of significant effects (logistic regression, p<0.05 for both coefficient Wald test and model likelihood ratio test). **(C-D).** Mean firing rate across units in each distance quartile during deviant trials for monkey B (left) and W (right). Responses closest to the fMRI cluster peak are in the top panel. Shaded regions indicate when fractal presentation occurs. Activity near the cluster peak shows a more phasic response with highest activity at the sequence start, while responses in more distant neurons show less modulation with image and sequence timing. Traces show mean ± SEM.

## Supplementary Tables

**Supplementary Table 1.** Activation coordinates from the human RD>NI model.

| **Contrast location** | **BA** | **Extent (voxels)** | **x** | **y** | **z** | **Peak t-val.** |
|---|---|---|---|---|---|---|
| Left LPFC (IFG, pars opercularis) | 44 | 2644 | -36 | 12 | 22 | 4.71 |
| Supplementary motor area/pre-SMA | 6 |  | 2 | 16 | 58 | 5.13 |
| Middle cingulate gyrus | 31 |  | -16 | -16 | 36 | 4.70 |
| Middle frontal gyrus | 9 |  | -26 | 8 | 34 | 4.45 |
| Precentral gyrus | 8 |  | -46 | 4 | 38 | 4.35 |
| Right supplementary motor area | 6 |  | 12 | 2 | 60 | 3.97 |
| Paracentral lobule | 4 |  | -6 | -18 | 64 | 3.95 |
| Middle cingulate gyrus | 32 |  | -10 | 12 | 44 | 3.55 |
| Precentral gyrus | 6 |  | -44 | -2 | 54 | 3.22 |
| Middle frontal gyrus | 9 |  | -38 | 22 | 38 | 2.97 |
| Right middle cingulate gyrus | 32 |  | 12 | 14 | 42 | 2.89 |
| Paracentral lobule | 5/4 |  | -10 | -36 | 68 | 2.86 |
| Right LPFC (IFG, pars triangularis) | 45 | 1327 | 40 | 18 | 20 | 4.08 |
| Precentral gyrus | 6 |  | 46 | 4 | 28 | 3.85 |
| IFG, pars triangularis | 45 |  | 54 | 30 | 18 | 3.54 |
| Precentral gyrus | 6 |  | 54 | 4 | 46 | 3.50 |
| Middle frontal gyrus | 6 |  | 38 | 0 | 52 | 3.14 |
| Middle frontal gyrus | 9 |  | 32 | 24 | 34 | 2.84 |

*All local maxima ≥16mm apart; height threshold p < 0.005 uncorrected, family-wise error cluster-correction extent k = 1327 voxels, df = 27. Coordinates in MNI.*

**Supplementary Table 2.** Activation coordinates from the monkey RD>NI model.

| Contrast location | BA | Extent (voxels) | x | y | z | Peak t-val. |
|---|---|---|---|---|---|---|
| Medial prefrontal cortex | 10mr | 197 | 0.5 | 47.5 | 17.5 | 6.24 |
|  |  |  | -1 | 45.5 | 16 | 4.23 |
|  |  |  | -3 | 46 | 15 | 3.65 |
|  |  |  | -1.5 | 47.5 | 15 | 3.60 |
| Dorsolateral prefrontal cortex | 46d | 206 | 9.5 | 35 | 22 | 5.44 |
|  |  |  | 12.5 | 34 | 23.5 | 4.58 |
|  |  |  | 12 | 36 | 23.5 | 3.81 |
|  |  |  | 14 | 36.5 | 26 | 3.29 |
| Caudate (Cd) |  | 183 | 13 | 15.5 | 7 | 5.18 |
| Putamen (Pu) |  |  | 14.5 | 14.5 | 8 | 3.92 |
|  |  |  | 14.5 | 12 | 9 | 3.78 |
| Caudate (Cd) |  |  | 14 | 13.5 | 6 | 3.63 |
| Auditory cortex |  | 462 | -20 | 14 | 9 | 5.09 |
|  |  |  | -25 | 14.5 | 12.5 | 4.68 |
| Caudate (Cd) |  |  | -15 | 11.5 | 7 | 4.33 |
| Auditory cortex |  |  | -22.5 | 13.5 | 10 | 4.13 |
| Caudate (Cd) |  | 411 | 11 | 21 | 24 | 4.44 |
| Somatosensory cortex | 3a/b |  | 12 | 15 | 25 | 4.00 |
| Anterior cingulate cortex | 24c |  | 9 | 19 | 26.5 | 3.50 |
| Dorsal premotor cortex (PMd) | 6 |  | 13 | 22.5 | 24.5 | 3.40 |
| Cerebellum |  | 221 | 11.5 | -9.5 | 5.5 | 4.43 |
|  |  |  | 13 | -11 | 4.5 | 3.73 |
|  |  |  | 9.5 | -6.5 | 5.5 | 2.99 |
|  |  |  | 7.5 | -4.5 | 6 | 2.95 |

*All local maxima ≥2mm apart; height threshold p < 0.005 uncorrected, family-wise error cluster-correction extent k = 183 voxels, df = 21. Coordinates in 112RM-SL.*

**Supplementary Table 3.**
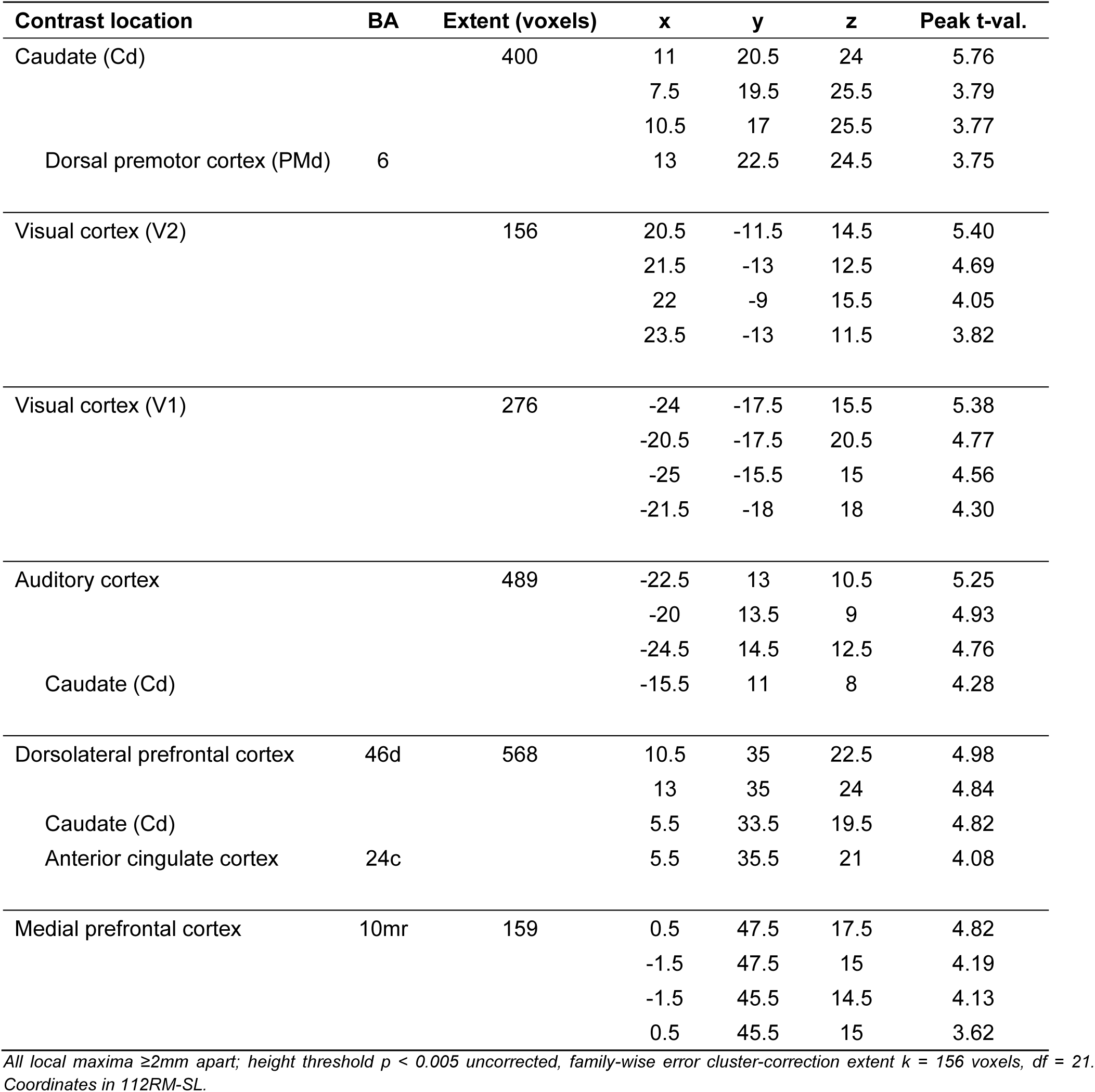
Monkey Rp46f-to-Monkey ISC correlation coordinates.

| **Contrast location** | **BA** | **Extent (voxels)** | **x** | **y** | **z** | **Peak t-val.** |
|---|---|---|---|---|---|---|
| Caudate (Cd) |  | 400 | 11 | 20.5 | 24 | 5.76 |
|  |  |  | 7.5 | 19.5 | 25.5 | 3.79 |
|  |  |  | 10.5 | 17 | 25.5 | 3.77 |
| Dorsal premotor cortex (PMd) | 6 |  | 13 | 22.5 | 24.5 | 3.75 |
| Visual cortex (V2) |  | 156 | 20.5 | -11.5 | 14.5 | 5.40 |
|  |  |  | 21.5 | -13 | 12.5 | 4.69 |
|  |  |  | 22 | -9 | 15.5 | 4.05 |
|  |  |  | 23.5 | -13 | 11.5 | 3.82 |
| Visual cortex (V1) |  | 276 | -24 | -17.5 | 15.5 | 5.38 |
|  |  |  | -20.5 | -17.5 | 20.5 | 4.77 |
|  |  |  | -25 | -15.5 | 15 | 4.56 |
|  |  |  | -21.5 | -18 | 18 | 4.30 |
| Auditory cortex |  | 489 | -22.5 | 13 | 10.5 | 5.25 |
|  |  |  | -20 | 13.5 | 9 | 4.93 |
|  |  |  | -24.5 | 14.5 | 12.5 | 4.76 |
| Caudate (Cd) |  |  | -15.5 | 11 | 8 | 4.28 |
| Dorsolateral prefrontal cortex | 46d | 568 | 10.5 | 35 | 22.5 | 4.98 |
|  |  |  | 13 | 35 | 24 | 4.84 |
| Caudate (Cd) |  |  | 5.5 | 33.5 | 19.5 | 4.82 |
| Anterior cingulate cortex | 24c |  | 5.5 | 35.5 | 21 | 4.08 |
| Medial prefrontal cortex | 10mr | 159 | 0.5 | 47.5 | 17.5 | 4.82 |
|  |  |  | -1.5 | 47.5 | 15 | 4.19 |
|  |  |  | -1.5 | 45.5 | 14.5 | 4.13 |
|  |  |  | 0.5 | 45.5 | 15 | 3.62 |

*All local maxima ≥2mm apart; height threshold p < 0.005 uncorrected, family-wise error cluster-correction extent k = 156 voxels, df = 21. Coordinates in 112RM-SL.*

**Supplementary Table 4.** Human right LPFC-to-Human ISC correlation coordinates.

| Contrast location | BA | Extent (voxels) | x | y | z | Peak t-val. |
|---|---|---|---|---|---|---|
| Right LPFC (IFG, pars triangularis) | 45 | 18032 | 46 | 22 | 22 | 10.55 |
| Right cuneus | 19 |  | 14 | -70 | 34 | 9.92 |
| Right LPFC (IFG, pars triangularis) | 44 |  | 42 | 26 | 10 | 7.13 |
| Right LPFC (MFG) | 46 |  | 40 | 42 | 16 | 7.11 |
| Left lingual gyrus | 17 |  | 2 | -80 | 2 | 6.95 |
| Left LPFC (IFG, pars triangularis) | 44 |  | -44 | 16 | 26 | 6.88 |
| Right LPFC (IFG, pars opercularis) | 44 |  | 52 | 10 | 20 | 6.66 |
| Right putamen | 44 |  | 32 | 14 | 4 | 6.57 |
| Right middle cingulate cortex | 32 |  | 10 | 18 | 32 | 6.08 |
| Right insula | 47 |  | 34 | 26 | -2 | 6.07 |
| Left insula | 45 |  | -34 | 22 | 6 | 6.05 |
| Left superior occipital gyrus | 7 |  | -14 | -72 | 28 | 6.03 |
| Right LPFC (SFG) | 6 |  | 24 | 12 | 60 | 6.02 |
| Left fusiform gyrus | 35-36 | 1237 | -34 | -20 | -30 | 8.96 |
| Left inferior temporal gyrus | 20 |  | -48 | -30 | -28 | 7.45 |
|  |  |  | -46 | -14 | -38 | 7.15 |
| Right middle temporal gyrus | 21 | 4131 | 68 | -46 | 2 | 7.80 |
| Right superior temporal gyrus | 22 |  | 52 | -8 | -12 | 7.50 |
| Right middle temporal gyrus | 21 |  | 50 | -22 | -12 | 7.50 |
| Right inferior temporal gyrus | 21 |  | 64 | -52 | -10 | 6.12 |
| Right superior temporal gyrus | 21 |  | 46 | -42 | 18 | 5.69 |
| Right supramarginal gyrus | 40 |  | 64 | -24 | 40 | 5.47 |
| Right superior temporal gyrus | 41-42-52 |  | 54 | -22 | 12 | 4.97 |
| Right inferior temporal gyrus | 20 |  | 40 | -8 | -44 | 4.85 |
| Right supramarginal gyrus | 22 |  | 64 | -40 | 26 | 4.73 |
| Right fusiform gyrus | 20 |  | 36 | -2 | -32 | 4.72 |
| Right postcentral gyrus | 2 |  | 34 | -32 | 40 | 4.64 |
| Left superior temporal gyrus | 22 | 1822 | -64 | -52 | 14 | 5.85 |
| Left inferior parietal lobule | 2 |  | -54 | -24 | 38 | 5.62 |
| Left superior temporal gyrus | 41-42-52 |  | -62 | -32 | 12 | 5.19 |
|  | 22 |  | -52 | -44 | 18 | 4.91 |
| Left postcentral gyrus | 40 |  | -50 | -30 | 50 | 4.70 |

*All local maxima ≥13mm apart; height threshold p < 0.005 uncorrected, family-wise error cluster-correction extent k = 1237 voxels, df = 27. Coordinates in MNI. Large, variable cluster sizes motivated listing different numbers of peaks across clusters.*

**Supplementary Table 5.** Monkey Rp46f-to-Human correlation coordinates.

| **Contrast location** | **BA** | **Extent (voxels)** | **x** | **y** | **z** | **Peak t-val.** |
|---|---|---|---|---|---|---|
| Right LPFC (IFG, pars triangularis) | 45 | 458 | 42 | 24 | 18 | 5.49 |
|  | 44 |  | 52 | 18 | 24 | 4.12 |
|  | 44 |  | 38 | 14 | 22 | 3.66 |
| Right superior temporal gyrus | 22 | 302 | 64 | -38 | 16 | 4.98 |
|  | 22 |  | 56 | -38 | 14 | 3.75 |
|  | 22 |  | 48 | -40 | 16 | 3.74 |
| Right middle temporal gyrus | 21 | 292 | 56 | -38 | -8 | 4.55 |
|  | 21 |  | 50 | -28 | -8 | 3.91 |
| Supplementary motor area | 6 | 696 | -2 | 4 | 60 | 4.38 |
|  | 6 |  | 0 | 18 | 58 | 4.35 |
|  | 6 |  | 6 | 8 | 54 | 4.24 |
| Left superior temporal gyrus | 22 | 390 | -64 | -48 | 16 | 4.36 |
|  | 22 |  | -68 | -40 | 14 | 4.31 |
|  | 22 |  | -68 | -32 | 12 | 4 |
| Left LPFC (IFG, pars opercularis) | 44 | 239 | -40 | 14 | 22 | 3.93 |
| Left precentral gyrus |  |  | -36 | 4 | 38 | 3.74 |
| Left LPFC (IFG, pars triangularis) |  |  | -38 | 22 | 26 | 3.49 |

*All local maxima ≥8mm apart; height threshold p < 0.005 uncorrected, family-wise error cluster-correction extent k = 239, df = 27. Coordinates in MNI.*

**Supplementary Table 6.** Human right LPFC-to-Monkey ISC correlation coordinates.

| Contrast location | BA | Extent (voxels) | x | y | z | Peak t-val. |
|---|---|---|---|---|---|---|
| Visual cortex (V2) |  | 197 | -7.5 | -14.5 | 30 | 5.58 |
| V6 |  |  | -3 | -10.5 | 29 | 5.41 |
| V2 |  |  | -5 | -12.5 | 29 | 4.29 |
|  |  |  | -7.5 | -12 | 29.5 | 4.17 |
| Visual cortex (V2) |  | 232 | 2.5 | -12.5 | 30 | 5.22 |
| V3 |  |  | 4 | -14 | 33 | 4.96 |
| V2 |  |  | 2.5 | -12.5 | 34 | 4.61 |
|  |  |  | 4.5 | -14 | 30.5 | 4.45 |
| Caudate (Cd) |  | 184 | 6 | 33 | 18.5 | 4.54 |
| Dorsolateral prefrontal cortex | 46 |  | 7 | 36 | 20.5 | 3.52 |
| Visual cortex (V2) |  | 189 | 4 | -18 | 17 | 4.42 |
| V1 |  |  | 3.5 | -17 | 19 | 4.28 |
| V2 |  |  | 8 | -16.5 | 18.5 | 4.11 |
|  |  |  | 5.5 | -16 | 20 | 3.43 |

*All local maxima ≥2mm apart; height threshold p < 0.005 uncorrected, family-wise error cluster-correction extent k = 184 voxels, df = 21. Coordinates in 112RM-SL.*

**Supplementary Table 7.** Summary of task variable coding in LPFC units.

| **Subject** | **Time Bin** | *Ordinal Position* | *Sequence Category* | *Stimulus Identity (deviant)* | *Stimulus Identity (habituation)* |
|---|---|---|---|---|---|
| Monkey B<br>(n = 469) | *ImgOn* | 197 | -7.5 | -14.5 | 30 |
|  | *ImgOff* | 232 | 2.5 | -12.5 | 30 |
| Monkey W<br>(n = 418) | *ImgOn* | 184 | 6 | 33 | 18.5 |
|  | *ImgOff* | 189 | 4 | -18 | 17 |

*Ordinal Position and Sequence Category: 2-way ANOVA with an interaction term, p < 0.05. Deviant Type, Stimulus Identity (habituation trials), and Stimulus Identity (deviant trials): separate one-way ANOVAs, p < 0.05.*

**Supplementary Table 8.** Monkey-Specific fMRI Peak Coordinates.

| **Subject** | **RD>NI Cluster Peak** |
|---|---|
| Monkey B | 13.0, 39.8, 28.3 |
| Monkey W | -20.5, 28.8, 25 |

*Individual monkey contrasts in their native space, thresholded at p < 0.05 (uncorrected). Coordinates correspond to millimeters distance from ear-bar-zero each monkey’s stereotactic space (xyz = ML, AP, depth).*

**Supplementary Table 9.** Logistic regression of task variable coding frequency as a function of cluster distance for Monkey B.

|  | ***Monkey B, ImgOn*** |  |  | ***Monkey B, ImgOff*** |  |  |
|---|---|---|---|---|---|---|
|  | *β* | *p<sub>coef</sub>* | *p<sub>model</sub>* | *β* | *p<sub>coef</sub>* | *p<sub>model</sub>* |
| *Pos1* | **-0.67** | **4.5E-10** | **7.2E-12** | **-1.14** | **1.3E-11** | **2.5E-16** |
| *Pos2* | 0.75 | 0.14 | 0.10 | 0.10 | 0.89 | 0.89 |
| *Pos3* | -0.23 | 0.40 | 0.39 | -0.10 | 0.72 | 0.71 |
| *Pos4* | -0.083 | 0.76 | 0.76 | -0.50 | 0.086 | 0.071 |
| *Hab>Dev* | 0.12 | 0.38 | 0.38 | 0.26 | 0.10 | 0.093 |
| *Dev>Hab* | 0.15 | 0.16 | 0.16 | -0.059 | 0.58 | 0.58 |
| *NI>RD* | -7.8E-3 | 0.96 | 0.96 | 0.069 | 0.65 | 0.65 |
| *RD>NI* | 0.29 | 0.065 | 0.061 | 0.067 | 0.61 | 0.61 |

*Separate logistic models were run for each task variable, subtype, and time window. Negative β indicates that the variable and subtype preference were more common closer to the fMRI cluster peak. Variables with p < 0.05 for both coefficient (Wald test) and model (likelihood ratio test) are shown in bold.*

**Supplementary Table 10.**
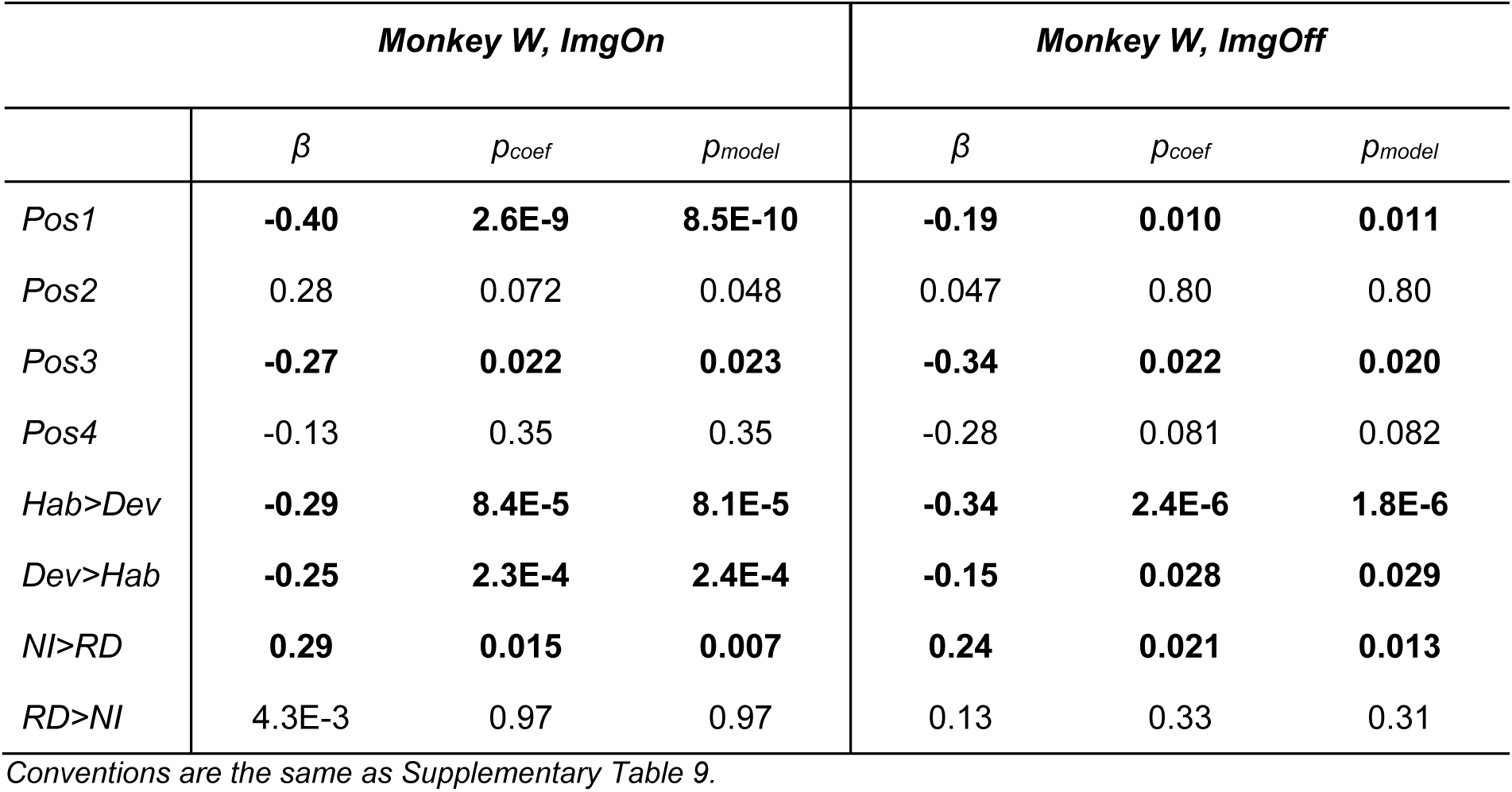
Logistic regression of task variable coding frequency as a function of cluster distance for Monkey W.

| ***Monkey W, ImgOn*** |  |  |  | ***Monkey W, ImgOff*** |  |  |
|---|---|---|---|---|---|---|
|  | *β* | *p<sub>coef</sub>* | *p<sub>model</sub>* | *β* | *p<sub>coef</sub>* | *p<sub>model</sub>* |
| *Pos1* | **-0.40** | **2.6E-9** | **8.5E-10** | **-0.19** | **0.010** | **0.011** |
| *Pos2* | 0.28 | 0.072 | 0.048 | 0.047 | 0.80 | 0.80 |
| *Pos3* | **-0.27** | **0.022** | **0.023** | **-0.34** | **0.022** | **0.020** |
| *Pos4* | -0.13 | 0.35 | 0.35 | -0.28 | 0.081 | 0.082 |
| *Hab>Dev* | **-0.29** | **8.4E-5** | **8.1E-5** | **-0.34** | **2.4E-6** | **1.8E-6** |
| *Dev>Hab* | **-0.25** | **2.3E-4** | **2.4E-4** | **-0.15** | **0.028** | **0.029** |
| *NI>RD* | **0.29** | **0.015** | **0.007** | **0.24** | **0.021** | **0.013** |
| *RD>NI* | 4.3E-3 | 0.97 | 0.97 | 0.13 | 0.33 | 0.31 |

*Conventions are the same as Supplementary Table 9.*

